# An expandable FLP-ON::TIR1 system for precise spatiotemporal protein degradation in *C. elegans*

**DOI:** 10.1101/2022.10.14.512315

**Authors:** Yutong Xiao, Callista Yee, Michael A. Q. Martinez, Chris Z. Zhao, Wan Zhang, Kang Shen, David Q. Matus, Christopher Hammell

## Abstract

The auxin-inducible degradation system has been widely adopted in the *C. elegans* research community for its ability to empirically control the spatiotemporal expression of target proteins. This system can efficiently degrade <u>*a*</u>uxin-<u>*i*</u>nducible <u>*d*</u>egron (AID)-tagged proteins via the expression of a ligand-activatable _At_TIR1 protein derived from *A. thaliana* that adapts target proteins to the endogenous *C. elegans* proteosome. While broad expression of _At_TIR1 using strong, ubiquitous promoters can lead to rapid degradation of AID-tagged proteins, cell type-specific expression of _At_TIR1 using spatially restricted promoters often results in less efficient target protein degradation. To circumvent this limitation, we have developed a FLP/FRT_3_-based system that functions to reanimate a dormant, high-powered promoter that can drive sufficient _At_TIR1expression in a cell type-specific manner. We benchmark the utility of this system by generating a number of tissue specific FLP-ON::TIR1 drivers to reveal genetically separable cell type-specific phenotypes for several target proteins. We also demonstrate that the FLP-ON::TIR1 system is compatible with enhanced degron epitopes. Finally, we provide an expandable toolkit utilizing the basic FLP-ON::TIR1 system that can be adapted to drive optimized _At_TIR1expression in any tissue or cell type of interest.

## Introduction

The capability to regulate the expression of transgenes and endogenous protein activity has revolutionized the study of biology in multicellular genetic model organisms. The invariant cell lineage of *C. elegans* and genetic tractability of the system provide a unique opportunity to study how aspects of gene regulation and cell biology directly correlate with the establishment of individual cell fates. A number of molecular techniques have been developed to perturb gene expression including RNA interference (RNAi) in sensitized backgrounds (Simmer et al. 2002; Watts et al. 2020) and controlled elimination of gene activity using targeted DNA recombination through Cre/lox or FLP/FRT techniques (Driesschaert et al. 2021; Nance and Frokjaer-Jensen 2019). While both methods can be adapted for tissue-specific inactivation of target gene expression, they are limited in their utility by the persistence of protein expression following the destruction of the targeted mRNA or deletion of the genomic loci. These systems have been complemented by a number of hybrid approaches that directly target proteins of interest for degradation. These include ZF1-tagging systems (Armenti et al. 2014), GFP nanobody approaches (Wang et al. 2017), and the auxin inducible degradation (AID) system (Zhang et al. 2015). Each of these strategies require the addition of a specific epitope or peptide to the target gene as well as the expression of one of more transgenes that recognize and adapt the target to the protein turnover machinery.

Of these systems, the AID system has been widely adopted for its ability to rapidly and specifically induce degradation of auxin-inducible degron (AID)-tagged proteins. The ease of editing the *C. elegans* genome using CRISPR/Cas9 (Dickinson et al. 2015) allows a target gene of interest to be efficiently tagged with a small 44 amino acid epitope (AID) that can be recognized by _At_TIR1, a heterologously expressed protein derived from *A. thaliana*. _At_TIR1 serves as a substrate-recognition component that tethers target proteins to the endogenous *C. elegans* SKP1-CUL1-F-box (SCF) E3 ubiquitin ligase complex in the presence of a cuticle-permeable, plant hormone called indole-3-acetic acid (IAA or auxin). Therefore, exquisite control of the temporal aspects of degradation can be experimentally controlled as AID-tagged target proteins can only be degraded after exogenous auxin exposure.

The original AID system has been improved upon iteratively, which has led to increased functionality in *C. elegans-specific* contexts. For example, an eggshell-permeable analog of IAA, acetoxymethyl indole-3-acetic acid (IAA-AM), was synthesized to target AID-tagged proteins for degradation in developing embryos (Negishi et al. 2019). For targeted degradation in microfluidics systems or liquid cultures, a synthetic analog that exhibits increased solubility in aqueous solutions, 1-naphthaleneacetic acid (NAA), and its potassium salt K-NAA, can be employed (Martinez et al. 2020; Martinez and Matus 2020). Mutation of amino acids within the binding pocket (phenylalanine 79 to glycine (F79G)) that alter the ligand specificity of _At_TIR1 to a modified IAA analog (5-phenyl-indole-3-acetic acid (5-Ph-IAA)) leads to a dramatic reduction of ligand-independent _At_TIR1 activity and also improves the efficacy of targeted degradation (Hills-Muckey et al. 2022; Negishi et al. 2022). Finally, an expanded AID system toolkit has been developed that offers a set of single-copy, tissue-specific _AT_TIR1-expressing strains to adapt the AID/_At_TIR1 system for cell type-specific degradation in *C. elegans* (Ashley et al. 2021a).

While the AID/_At_TIR1 technology is a powerful tool to control targeted protein degradation, several reports indicate that cell type-specific application of the system may still need further optimization. For instance, depletion of multiple AID-tagged proteins using tissue-specific _At_TIR1 drivers fails to recapitulate the genetic null phenotypes. This is especially true when AID-targets are potent transcription factors (Patel and Hobert 2017), abundant structural components (Riga et al. 2021) and dosage-dependent regulators of cellular or developmental activities (Smith et al. 2022; van der et al. 2020). Given that the AID system relies only on an AID-tagged target protein, _At_TIR1 expression, and auxin (which is typically added at saturating concentrations), we reasoned that incomplete degradation of target proteins, in many contexts, may be due to insufficient or developmentally dynamic expression of _At_TIR1 expression from some tissue specific promoters.

Here, we engineer a hybrid transgenic system that programs cell type-specific FLP (flippase) activity to reanimate a dormant, high-powered, universal promoter to drive optimized _AT_TIR1(F79G) expression in a cell type-specific fashion. This composite system takes advantage of the large knowledgebase of previous studies of tissue-specific promoters and enables high-activity, _At_TIR1-dependent degradation to be achieved in specific cell types without a dependency on specific promoter strength. We benchmark the utility of this system by generating tissue specific FLP-ON::TIR1 drivers and use them to reveal genetically separable, cell type-specific phenotypes for multiple important regulators of differential cell identity. Finally, we build an expandable toolkit utilizing the basic FLP-ON::TIR1 system and demonstrate how it can be easily adapted to drive _AT_TIR1(F79G) expression in desired tissues or cell types.

## Results

### AID::GFP degradation is limited by TIR1 expression levels when driven by some tissue-specific promoters

Previous kinetic experiments from our laboratory and others have demonstrated that degradation of *eft-3p*::AID::GFP (Zhang et al. 2015) is highly efficient in strains that express _At_TIR1 using strong, ubiquitous promoters, such as *rpl-28* (Hills-Muckey et al. 2022) (Fig. 1A, B’). In our effort to harness this capability in a cell type-specific manner, we drove an identical _At_TIR1::P2A::mCh::HIS-11 transgene in the vulval precursor cells (VPC) using a well-characterized variant of the *lin-31* promoter (Tan et al. 1998) (Fig. 1A). Surprisingly, we found that the kinetics of *lin-31p*:: _At_TIR1-dependent depletion of the *eft-3p*::AID::GFP in the VPCs was significantly slower than what is observed in these same cells in animals expressing the ubiquitous *rpl-28p*::_At_TIR1 construct (Fig. 1B”). Specifically, the ubiquitous _At_TIR1 driver depleted greater than 50% of the AID-tagged GFP protein within the first 30 minutes of auxin addition (1 mM K-NAA) while _At_TIR1 expressed from the *lin-31* promoter resulted in only 5% of target protein degradation (Fig. 1F). We hypothesized that this difference in activity may result from differences in the expression levels of _At_TIR1 in each context. To query this, we measured the expression levels of a co-translated and proteolytically cleaved mCh::HIS-11 reporter that is derived from each _At_TIR1::P2A::mCh::HIS-11 transgene. (Ahier and Jarriault 2014). Based on quantification of mCh::HIS-11 expression, we estimated that _At_TIR1 expression levels derived from the *lin-31* prompter are approximately 2.5-fold lower than the levels expressed in VPCs from the *rpl-28* promoter (Fig. 1C-D), indicating insufficient degradation of *eft-3p*::AID::GFP likely results from the lower level of _At_TIR1 expressed by the *lin-31* promoter.

**Figure 1.**
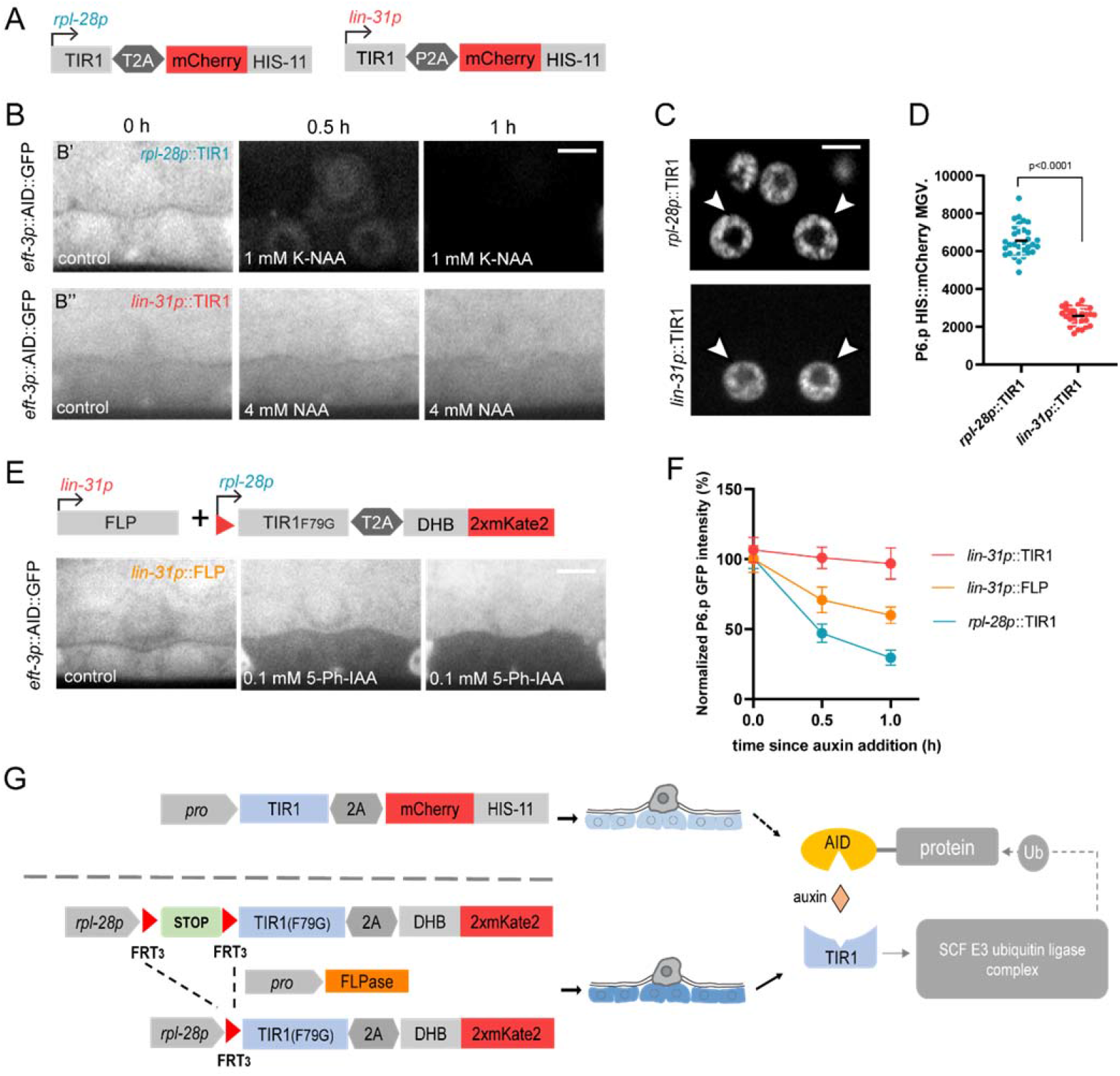
*lin-31p*::FLP mediated recombination can efficiently degrade AID::GFP specifically in the VPCs. (A) Schematic of *rpl-28p*::TIR1::T2A::mCh::HIS-11 and *lin-31p*::TIR1::P2A::mCh::HIS-11 constructs. (B) Micrographs depicting *eft-3p*::AID::GFP depletion following auxin addition using *rpl-28p*::TIR1 and *lin-31p*::TIR1. (C) Micrographs depicting mCh::HIS-11 expression by *rpl-28p* and *lin-31p*. White arrow heads indicate mCh::HIS-11 in P6.p. (D) Quantification of mCh::HIS-11 MGV (mean gray value) in P6.p. Student’s *t*-test compared between *rpl-28p*::TIR1 and *lin-31p::TIR1*. n≥30 animals per treatment. (E) Schematic of *lin-31p::FLP* mediated *rpl-28p*::>::_At_TIR1(F79G)::T2A::DHB::2xmK2 recombination. Micrographs depicting FLP-ON^lin-31^::TIR1 specifically degraded *eft-3p*::AID::GFP in P6.p. (F) Quantification of normalized *eft-3p*::AID::GFP MGV in P6.p., each data point represents the average intensity in P6.p. for the given condition and timepoint. Error bars: mean ± SD; n: ≥30 animals. (G) Schematic figure of comparison between *promoter*::TIR1 and FLP-ON::TIR1 system. Scale bars: 5 μm.

### Reanimation of an inert transgene using tissue-specific FLP activity enables maximal TIR1 expression in specific cell types

A common approach to drive conditional gene expression is to use targeted DNA recombination (Nance and Frokjaer-Jensen 2019; Nonet 2020) to generate a highly active transgene from a parental construct that includes a stop cassette that prevents transcriptional read-through of the reporter. In these systems, cell type-specific expression of a recombinase such as FLP or Cre results in the excision of the stop cassette, restoring efficient transgene expression only in cell types harboring recombinase activity (Davis et al. 2008; Voutev and Hubbard 2008). We elected to optimize the FLP/FRT_3_ recombination system in this context as this FLP/FRT_3_ combination has previously been demonstrated to work efficiently in *C. elegans* (Munoz-Jimenez et al. 2017). We then generated a *rpl-28*p::>STOP>::_At_TIR1(F79G)::T2A::DHB::2xmK2 construct that harbors a >STOP> cassette (“>” denotes the FRT_3_ site) immediately upstream of the _At_TIR1(F79G)::T2A::DHB::2xmK2 sequence (Martinez et al. 2022). Next, using CRISPR/Cas9 genome engineering we inserted this construct into the *C. elegans* genome at a defined safe harbor site on chromosome II, corresponding to the ttTi5605 MosSCI insertion site (Frokjaer-Jensen et al. 2008). Consistent with the hypothesis that the >STOP> cassette prevents normal expression of the downstream encoded transgene, expression of the co-transcribed CDK activity sensor, DHB::2xmK2 (Adikes et al. 2020) was not detected in transgenic animals.

To determine if this transgene can be re-animated in a cell type-specific manner, we generated an expression construct that utilizes the same *lin-31* promoter used above to drive a variant of FLP (D5; aspartic acid to glycine mutation at aa 5) that exhibits elevated flippase activity (Nern et al. 2011) and a H2B::2xmT2 reporter. This construct was then inserted via CRISPR into a second safe harbor site on chromosome I, corresponding to the MosSCI site ttTi4348 (Philip et al. 2019). We then crossed each of these transgenes into the same genetic background (Fig. 1E). We found that 100% (n = 40) of late L2 larvae harboring both *rpl-28p*::>STOP>::_At_TIR1 (F79G)::T2A::DHB::2xmK2 and *lin-31p::FLP* transgenes efficiently express DHB::2xmK2 in all VPCs (Fig. S1), indicating that the FLP-ON^lin-31^::TIR1 system can efficiently generate functional _At_TIR1-expressing transgenes. We next determined if reanimation of *rpl-28p::*>::_At_TIR1(F79G)::T2A::DHB::2xmK2 expression in VPCs could more efficiently deplete the same *eft-3p*::AID::GFP reporter used in the experiments outlined in Fig. 1A. Addition of FLP-ON^lin-31^::TIR1 animals to media containing 0.1 mM 5-Ph-IAA led to a rapid and specific reduction of AID::GFP expression in VPCs (Fig. 1E). Importantly, the kinetics of AID::GFP degradation in FLP-ON^lin-31^::TIR1 animals was 8-to-10-fold faster when compared to the kinetics exhibited in animals expressing _At_TIR1 from the *lin-31* promoter (Fig. 1F). Given that our FLP-ON^lin-31^::TIR1 system achieved more efficient AID::GFP degradation with faster kinetics in a defined lineage, the VPCs, we were optimistic that this improved system could ameliorate the limitations of *promoter*::_At_TIR1 constructs. Thus, our next step was to test this system by performing functional perturbations using genes that have well-characterized null phenotypes.

### The FLP-ON::TIR1 system can be used to dissect cell type-specific activities of LIN-12/Notch

We first aimed to benchmark this approach using a variety of tissue- and cell-specific FLP drivers in developmental contexts where phenotypes associated with cell type-specific FLP-ON::TIR1 activity could be compared to those associated with genetic null mutations of the same target. For example, Notch-mediated intercellular communication plays important roles in multiple cell fate specification events in *C. elegans* development. Phenotypic and anatomical descriptions of larvae harboring mutations in *lin-12*, one of the two *C. elegans* notch genes, indicates that LIN-12 activity is necessary for cell fate specification in both the somatic gonad and vulval precursor cells (Greenwald 1998; 2012). While roles for *lin-12* in both cell types are clear when scored in a variety of loss-of-function mutants, it is cumbersome to assign with certainty distinct cell-autonomous phenotypes for *lin-12*. Additionally, it is difficult to disentangle these phenotypes from downstream effects of cell transformation that alter adjacent cell fate in a non-cell autonomous manner. Traditional mosaic analysis of *lin-12* function relies upon the spontaneous somatic loss of a free duplication (chromosomal fragment) and can only be followed by linkage with a cell biological marker. Nevertheless, animals with mosaic *lin-12* expression were initially used for a series of landmark experiments to show cell-autonomous and non-autonomous roles of *lin-12* in somatic cells (Seydoux and Greenwald 1989). While this powerful approach led to the discovery of the regulatory logic that employs LIN-12 activity in these contexts, these experiments and other approaches using RNAi and laser ablation, lack the ability to precisely control temporal aspects of LIN-12 activity. This limitation may occlude investigations of potential sequential activities of LIN-12 in these processes.

LIN-12/Notch signaling has been extensively studied in the context of vulval cell fate induction (reviewed in Schindler and Sherwood 2013). During vulval development an extended set of epithelial cells (P3.p-P8.p) on the ventral surface of larvae originally possess the potential to become vulval cells. Under normal circumstances only the central P5.p-P7.p cells are induced to generate the mature vulval structure (Sternberg 2005; Yochem 2007). This induction is mediated by a morphogen signal, LIN-3/EGF, secreted from the anchor cell (AC) that induces P6.p to assume the 1° VPC fate through Ras activation. The adjacent P5.p and P7.p cells, which receive less LIN-3 signal that the central P6.p precursor, adopt the 2° cell fate which is enforced by LIN-12/Notch-mediated lateral inhibition. (Yoo et al. 2004). Dysregulation of either signaling pathway leads to abnormal vulval fate patterning.

To specifically degrade LIN-12 in the VPCs, we generated a FLP-ON^unc-62^::TIR1 transgene using a ~3 kb fragment of the *unc-62 p*romoter that enables FLP to be robustly expressed in developing VPCs (Jiang et al. 2009). Animals harboring this FLP transgene and the *rpl-28p*::>STOP>::_At_TIR1(F79G)::T2A::DHB::2xmK2 transgene express DHB::2xmK2 in the expected tissues (Fig. 2A-C). Synchronized L1 larvae were then cultured with 0.1 mM 5-Ph-IAA while vulval cell division patterns and LIN-12::mNG::AID expression were monitored throughout the L3 and early L4 stages of development. The experiment revealed a complete lack of detectable LIN-12::mNG::AID expression in the VPCs (Fig. 2A-C merged channel) that co-express DHB::2xmK2 indicating functionality of the FLP-ON^unc-62^::TIR1 system to trigger cell type-specific degradation. Consistent with an elimination of LIN-12 activity in these cells, FLP-ON^unc-62^::TIR1 animals exposed to 5-Ph-IAA also exhibited vulval cell fate specification errors (Fig. 2A-C merged channel).

**Figure 2.**
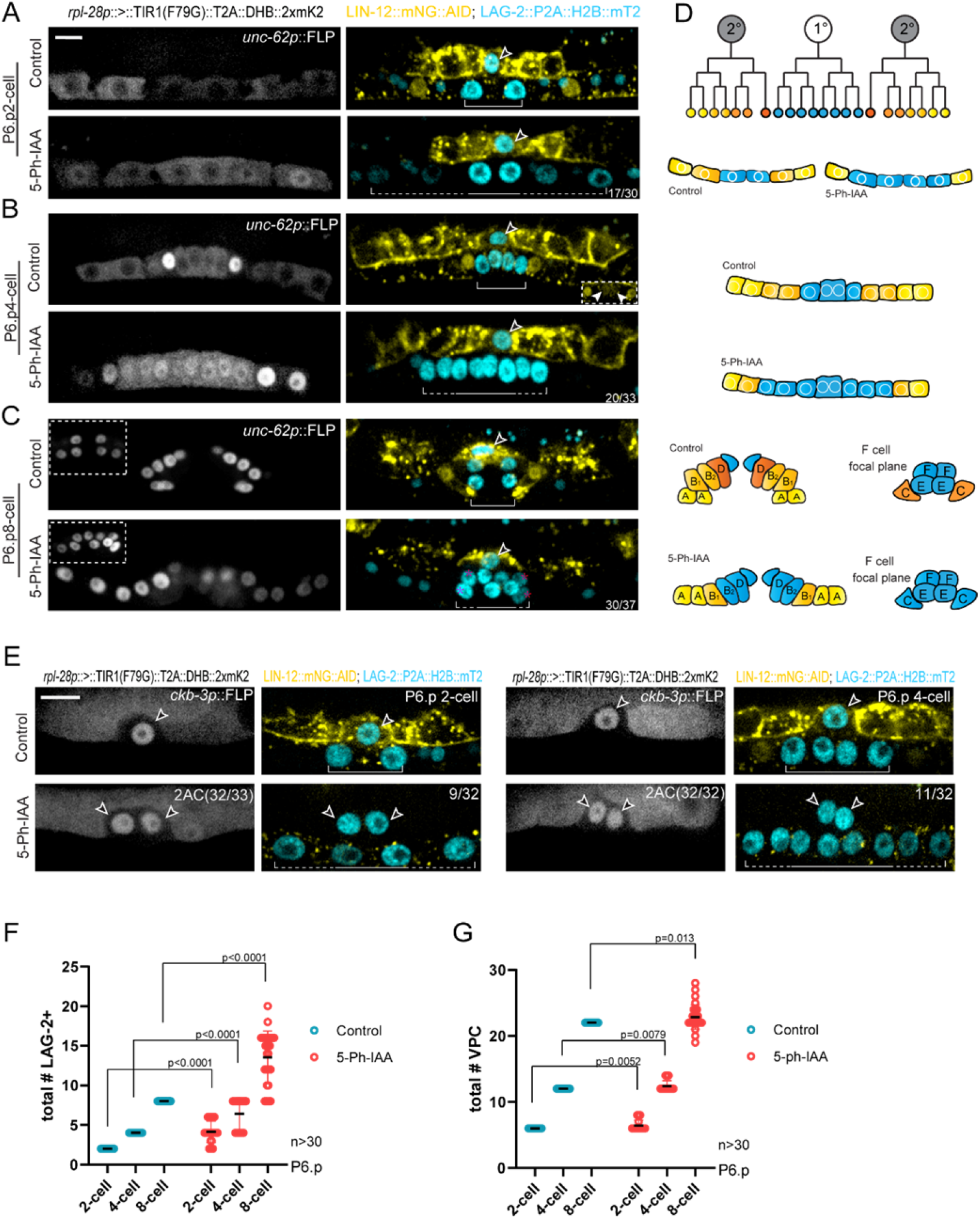
A FLP-ON::TIR1 system for dissecting LIN-12 functions in the VPCs and somatic gonad. (A-C) Representative images of normal and 5-Ph-IAA-mediated loss of LIN-12 phenotypes in VPCs via FLP-ON^unc-62^::TIR1 (normal 1° VPC pattern versus expanded 1° VPC) at P6.p 2-cell (A), P6.p4-cell (B), and P6.p 8-cell (C) developmental stage. Arrowheads indicates ACs. Solid bracket/line indicates normal P6.p LAG-2 expression pattern and dash line with bracket indicates expanded LAG-2 expression in P5.p and P7.p. Insets at P6.p 8-cell stage showing F cell focal plane. Pink asterisks indicates ectopic LAG-2 expression. (D) Schematic of vulva development and expanded primary fate after loss of LIN-12 in VPCs. P6.p expressing LAG-2 (blue) becomes 1° cell fate whereas P5.p and P7.p expressing LIN-12 (yellow and orange) become 2° cell fate. (E) Representative images of FLP-ON^ckb-3^::TIR1 induced loss of LIN-12 phenotypes in somatic gonad: 2AC 32/33 at P6.p 2-cell; 2AC 32/32 at P6.p 4-cell and extra ACs leads to expanded 1° vulval fate (9/32 at P6.p 2-cell; 11/32 at P6.p 4-cell), LIN-12::mNG::AID (yellow) merged with *lag-2p*::LAG-2::P2A::H2B::mT2 (blue). Arrow heads indicates ACs. Scale bar: 5 μm. (F, G) Column individual plots showing total LAG-2+ cells among VPCs (F) and total VPC number (G) based on DHB expression under control and 5-Ph-IAA treatment. P values calculated by Multiple t tests at each stage, n>30.

We scored the LIN-12::mNG::AID depletion phenotypes in developing vulval cells using two metrics. First, we quantified the number of vulval cells that ectopically express a 1° VPC cell fate by observing the expression of an endogenous transcriptional *lag-2* reporter (LAG-2::P2A::H2B::mT2) (Taylor N Medwig et al. 2022) at different vulval developmental stages. We observed that the number of *lag-2* (+) vulval cells with a 1° cell division pattern were significantly increased compared to untreated animals, indicating that cells that would normally adopt a 2° fate were transformed to a 1° cell fate when LIN-12 is depleted in VPCs (Fig. 2A-D, F). Second, we quantified the number of total vulval cells after auxin addition. In some 5-Ph-IAA-exposed animals, there were examples of extra vulval cells that may be derived from inappropriately induced P4.p, P8.p or ectopic division of C or D cells at certain stages (Fig. 2C, G).

In addition to the 1° cell fate expansion phenotype, we also observed aberrant cell-cycle states in transformed 2° cells. In vulval development, the innermost 2° vulval cell, the D cell, normally exits the cell-cycle one round of cell division earlier than the other vulval cells, resulting in the 22 terminally differentiated vulval cells in the L4 stage (Kiontke et al. 2007; Matus et al. 2014). This early cell-cycle exit can be easily distinguished by strong nuclear localization of DHB::2xmK2, our CDK activity sensor, shortly after mitotic exit (Adikes et al. 2020) (Fig. 2B). During the characterization of these *lin-12(-)* cell lineage transformations, we noticed that 1° fate expansion in 5-Ph-IAA-treated animals caused abnormal cell-cycle states in the D cell, as visualized by loss of nuclear DHB::2xmK2 (Fig. 2B). Additionally, in this LIN-12-degraded background, the neighboring A and B vulval cells appear to acquire the fates of their sister 2° cells, the C and D fates, with the “B” cell closest to the ectopic 1° cells adopting a D fate and exiting the cell-cycle into a CDK_low_ arrested state (Fig. 2B). Together, these data demonstrate that our FLP-ON^unc-62^::TIR1 system is able to degrade LIN-12/Notch in a specific tissue with expected loss-of-function phenotypes.

LIN-12/ Notch activity also plays a determinant role in the well-studied AC/VU decision within the somatic gonad (Iva S. Greenwald 1983). Initially, the inner-most proximal grand-daughters of the founder cells of the somatic gonad, Z1.ppp and Z4.aaa, have equal potential of becoming either a terminally differentiated AC or proliferative ventral uterine (VU) cell. The outcome of these mutually exclusive cell fate decisions is made through stochastic intracellular signaling events that are mediated by the receptor LIN-12/Notch and ligand LAG-2/Delta (Wilkinson et al. 1994). Cell lineage tracing of *lin-12(0)* mutants and laser ablation experiments indicate that *lin-12* activity is necessary for VU fate specification as both Z1.ppp and Z4.aaa adopt the default fate and become ACs in *lin-12(0)* mutants (Iva S. Greenwald 1983; Seydoux and Greenwald 1989).

We and others have tried unsuccessfully to reproduce *lin-12(0)* phenotypes by degrading LIN-12::mNG::AID in unspecified Z1.ppp and Z4.aaa (or the precursors of these cells) using gonad-specific promoters to express _At_TIR1. For example, our attempts to use the *ckb-3* promoter to drive _At_TIR1 expression in these cell types failed to achieve significant LIN-12::mNG::AID depletion in the somatic gonad and also failed to produce the associated cell transformation phenotypes (2 AC phenotype) (Fig. S2C) (Martinez et al. 2022; Taylor N Medwig et al. 2022). This inability to recapitulate the *lin-12(0)* phenotype is likely caused by an ineffective expression of _At_TIR1 in Z1.ppp and Z4.aaa or their precursors during the experimental time course. Indeed, the *ckb-3* promoter can drive transcriptional reporters in the gonad precursor cells Z1/Z4 and their decedents, but the level of promoter activity diminishes rapidly as the lineage progresses to more mature cell fates (Fig. S2A-B) (Benavidez et al. 2022; Shaffer and Greenwald 2022).

We therefore sought to adapt the FLP-ON::TIR1 system in this context to determine if we could effectively deplete LIN-12 in the somatic gonad. To accomplish this, we generated FLP-ON^ckb-3^::TIR1 using the *ckb-3* promoter (Benavidez et al. 2022; Shaffer and Greenwald 2022). This system was then combined with LIN-12::mNG::AID and the LAG-2::P2A::H2B::mT2 reporter transgene (Fig. 2E). The resulting strain specifically expressed DHB::2xmK2 in the somatic gonad as expected. After plating L1-synchronized animals onto solid media containing 0.1mM 5-Ph-IAA, we found that LIN-12::mNG::AID expression was specifically eliminated in the somatic gonad by the mid-L3 stage and that expression of LIN-12 persisted in developing VPCs (Fig. 2E). Using the LAG-2::P2A::H2B::mT2 reporter as a marker of AC fate (Wilkinson et al. 1994) and the CDK sensor to visualize cell-cycle state (Adikes et al. 2020), we observed 32/33 animals at the P6.p 2-cell stage and 33/33 animals at the P6.p 4-cell stage with two ACs arrested in a G0 state indicating that cells that would normally adopt the VU cell fate were efficiently transformed to the AC fate (Fig. 2E, merged channel). Thus, specifically degrading LIN-12::mNG::AID in the gonad using our FLP-ON^ckb-3^::TIR1 system recapitulates *lin-12(0)* phenotypes in Notch/Delta-mediated lateral inhibition during the AC/VU decision. This, combined with experiments utilizing the *unc-62* promoter in VPC cells, demonstrates that we can achieve cell type-specific degradation of LIN-12 in distinct cell types.

While depleting LIN-12::mNG::AID expression specifically in the developing ventral uterine cells, we also noticed that some VPCs exhibited a cell transformation phenotype (Fig. 2E merged channel). We hypothesized that this phenotype could arise from either of two causes. First, this may reflect an inappropriate expression of FLP activity in developing VPCs that was not anticipated from the previously described expression pattern of the *ckb-3* promoter, inappropriately causing Notch degradation in developing vulval cells. Alternatively, ectopic 1° fate transformation in adjacent VPCs could be triggered by a non-cell-autonomous mechanism mediated by excessive LIN-3/EGF activity derived from the ectopic induction of extra, LIN-12::mNG::AID depleted ACs/uterine cells. We reasoned that our FLP-ON::TIR1 system was perfectly suited to distinguish between these two outcomes. Consisted with the non-cell autonomous hypothesis, we failed to detect the presence of either H2B::2xmT2 or DHB::2xmK2 (Fig. S2B, D) in the VPCs, as expected, given that the Z1/Z4-specific *ckb-3p*::FLP activity was limited to the gonad. Second, LIN-12::mNG::AID in vulval cells following auxin treatment was comparable to untreated animals (Fig. 2C, merged channel), suggesting that the Notch-mediated lateral inhibition in the VPCs was not interrupted. Together, these results support the hypothesis that the ectopic 1° VPC fate transformation is caused by the excessive level of LIN-3/EGF secreted from additional ACs, which, in this case, is sufficient to overcome the normal level of lateral inhibition mediated by LIN-12/Notch activity in the VPCs. Importantly, this result highlights how the FLP-ON::TIR1 system can be used to unambiguously define cell type-specific functions for individual genes.

### The FLP-ON::TIR1 system identifies distinct functions of FOS-1A in the AC and other uterine cells

To further explore the efficacy of our FLP-ON::TIR1 system, we next sought to dissect the function of a key transcription factor, FOS-1, during somatic gonad development. The *fos-1* gene encodes the sole *C. elegans* homolog of the *fos* bZIP transcription factor family (Tatusov et al. 2003) and was initially identified as an essential regulatory component that controls the timing and organization of AC invasion through the basement membrane (BM)(Sherwood et al. 2005). A specific isoform of *fos-1, fos-1a*, is expressed exclusively in the developing gonad with early expression throughout the dorsal and ventral uterine cells. FOS-1A expression is also temporally controlled, increasing in expression in the AC following early AC specification (Sherwood et al. 2005; Medwig-Kinney et al. 2020). Mutations that specifically inactivate the expression of this isoform, *fos-1(ar105)*, lead to a fully penetrant AC invasion defect at the P6.p 4-cell stage, that prevents the required coupling of differentiating uterine cells with the developing vulva cells. These defects dramatically alter vulval morphogenesis and functionality (Sherwood et al. 2005).

To test if our FLP-ON::TIR1 system could robustly deplete FOS-1A and phenocopy *fos-1(ar105)*, we inserted mNG::AID coding sequences into the first exon of the endogenous *fos-1* locus by homology-directed repair using CRISPR/Cas9 genome engineering (Dickinson et al. 2015). We then combined this mNG::AID::FOS-1A allele with our gonad FLP-ON^ckb-3^::TIR1 system containing LAM-2::mNG as the BM reporter to score AC invasion defects following FOS-1A depletion. We exposed synchronized L1 larva to auxin for 24 hours to reach the mid-L3 and quantified the expression levels of FOS-1A in individual AC nuclei. In these conditions, FOS-1A was significantly degraded (Fig. 3B, >85% depletion in ACs) with a fully penetrant AC invasion defect such that 30/30 treated animals possessed intact LAM-2::mNG expression over P6.p-derived cells, indicative of a failure of the AC to invade (Fig. 3A, AC focal plane). These results demonstrate that our FLP-ON^ckb-3^::TIR1 system can recapitulate *fos-1(ar105)-specific* phenotypes.

**Figure 3.**
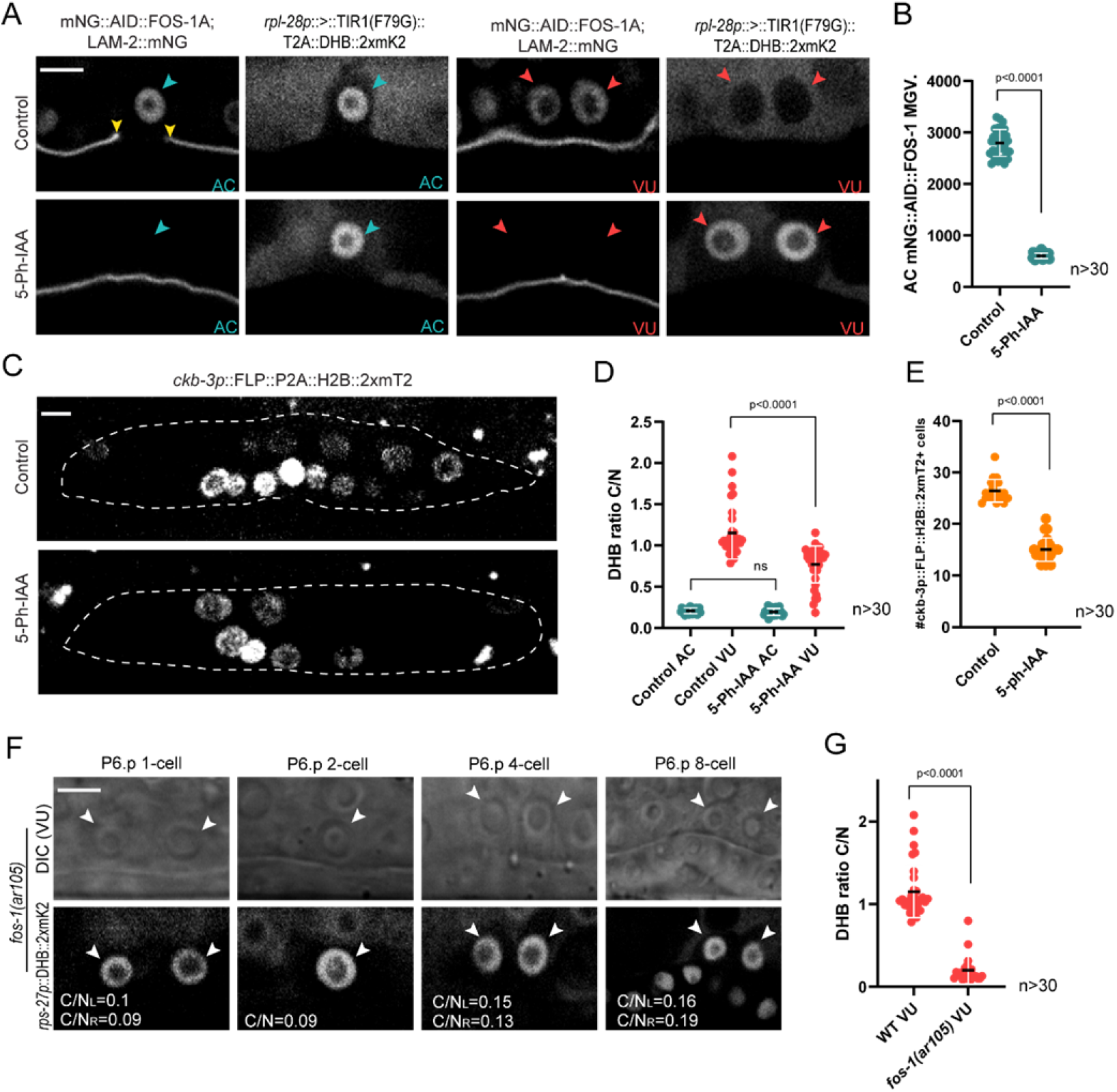
A FLP-ON::TIR1 system can identify distinct functions of FOS-1A in the AC and other uterine cells. (A) Representative images of mNG::AID::FOS-1A, basement membrane (BM, LAM-2::mNG), with FLP-ON^ckb-3^::TIR1 system to induce loss of FOS-1A in somatic gonad. AC focal plane labeled in blue: normal AC invasion versus invasion block. Open arrowheads indicate ACs. Yellow arrowheads indicate boundary of the breach on the BM. VU focal plane labeled in red: representative phenotypes of loss of FOS-1A in the VU, proliferating VUs versus cell-cycle arrested VUs. Solid arrowheads indicate VUs. (B) Quantification of mNG::AID-tagged FOS-1A MGV levels of individual AC nuclei, with FLP-ON^ckb-3^::TIR1 system (*n*≥30 animals per treatment, *P*<0.0001, statistical comparison was made by Student’s *t*-test between 5-Ph-IAA treated animals and control). (C) Somatic cells in the gonad with FLP-ON^ckb-3^::TIR1 induced loss of FOS-1A after 5-Ph-IAA treatment, (Maximum intensity projections of half z-stage, *ckb-3p::*FLP::P2A::H2B::2xmT2, collected by 63x objective, dash line indicates uterus,). (D) Quantification of C/N DHB::2xmK2 ratios in ACs and VUs (*n*≥30 animals per treatment, *P*<0.0001, statistical comparison was made by Student’s *t*-test between 5-Ph-IAA treated animals and control, n.s. not significant). (E) Number of somatic cells expressing *ckb-3p::*FLP::P2A::H2B::2xmT2 in the gonad with FLP-ON^ckb-3^::TIR1 induced loss of FOS-1A after 5-Ph-IAA treatment. (*n*≥30 animals, *P*<0.0001, statistical comparison was made by Student’s *t*-test between 5-Ph-IAA treated animals and control). (F) VU focal plane showing cell-cycle exit VU cells. Solid arrowheads indicate VUs. DHB ratios are demonstrated at left bottom corner. (G) Quantification of C/N DHB::2xmK2 ratios in wild type control and mutant VUs (*n*≥30 animals per condition, *P*<0.0001, statistical comparison was made by Student’s *t*-test between wild type and mutant).

During our analysis of AC-specific defects, we observed a novel, uncharacterized phenotype in adjacent uterine cells following FOS-1A depletion. Following auxin treatment, we observed an increase in cell size of uterine cells, suggesting a cell-cycle arrest phenotype in mNG::AID::FOS-1A-depleted animals. Using our cell-cycle sensor co-expressed in our FLP-ON^ckb-3^::TIR1 system, we quantified the CDK activity in these cells and found that the cell-cycle progression of DU/VU cells were dramatically affected following mNG::AID::FOS-1A degradation (Fig. 3A, VU focal plane), with significantly reduced mean DHB cytoplasm/nuclear ratio of 0.76 +/-0.22 (Fig. 3D). This is in striking contrast to wildtype VU and DU cells that are highly proliferative during this developmental window, with a variety of cell-cycle states distributed from S to G2 (Fig. 3D mean DHB ratio of 1.15+/-0.32) (Adikes et al. 2020). These data indicated that FOS-1A may be required for cell-cycle progression within the somatic gonad outside of the AC. To determine whether this is the case, we counted the number of cells expressing *ckb-3p*::FLP::P2A::2xmT2 in mid-L3 stage animals with and without auxin exposure. Consistent with our hypothesis, gonadal FOS-1A depletion led to half the wild-type number of somatic gonad cells (Fig. 3C, E). As this uterine cell-cycle arrest phenotype was not previously reported, we decided to monitor these same phenotypes in animals harboring the *fos-1(ar105)* allele. Following incorporation of a ubiquitously expressed CDK activity sensor, *rps-27p*::DHB::2xmK2 (Adikes et al. 2020), we observed large, arrested VU and DU cells in the *fos-1(ar105)* mutant animals, with a mean DHB ratio indicative of G0 arrest (Fig. 3F, G). Together, these data demonstrate the utility of lineage-specific targeted protein degradation and the ability to both phenocopy null allele phenotypes and uncover novel biology.

### The efficacy of AC-specific degradation of FOS-1A is improved by an alternative degron sequence, mIAA7

Given that FOS-1A depletion using a gonad-specific FLP-ON^ckb-3^::TIR1 system gave rise to phenotypes in multiple gonadal cell types, we next asked if we could more specifically target functions of FOS-1A that are unique to the invasive AC. To accomplish this, we fused an AC-specific 5 kb 5’ cis-regulatory element of the *lin-29* gene with FLP::P2A::H2B::2xmT2 (McClatchey et al. 2016) to generate FLP-ON^lin-29^::TIR1 and combined this system with the mNG::AID::FOS-1A allele described above. As expected, we detected *lin-29* promoter-dependent H2B::2xmT2 expression in the AC prior to invasion and did not detect expression in other somatic gonad cell types (Fig. S4). Following continuous auxin exposure from the L1 stages, we quantified a 58% depletion of FOS-1A in the AC at the P6.p 4-cell stage of development and found that this level of FOS-1A depletion leads to low penetrance of invasion defects (7/30 animals exhibited an intact BM) (Fig. 4A, D, E). These results suggest that robust AC-specific degradation may be more challenging using the FLP-ON^lin-29^::TIR1 system.

**Figure 4.**
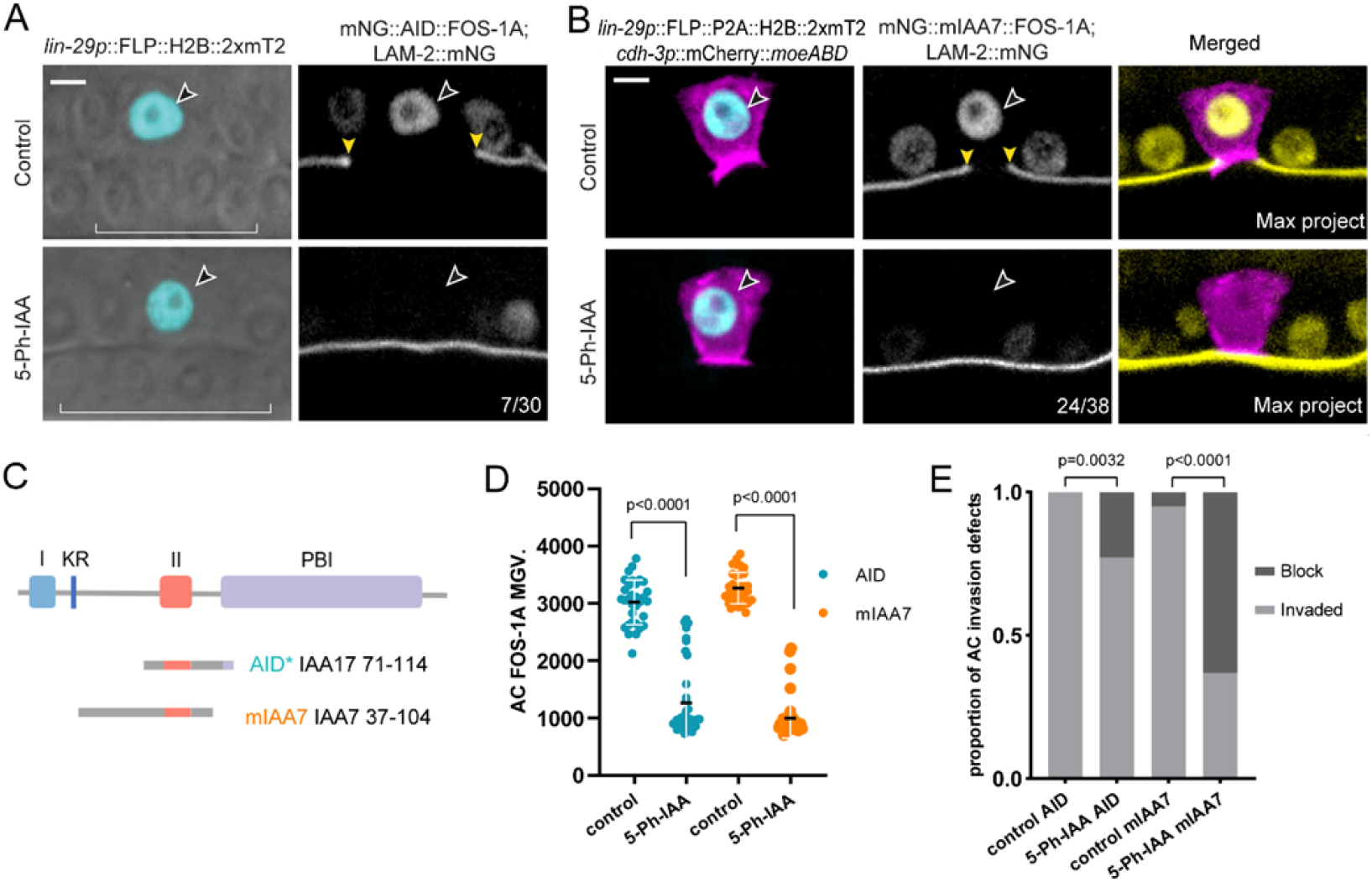
AC specific degradation of FOS-1A using AID and mIAA7 degrons. (A) Representative images of FLP-ON^lin-29^::TIR1 system (*lin-29p*::FLP::P2A::H2B::2xmT2), BM (LAM-2::mNG) with mNG::AID::FOS-1A. 5-Ph-IAA induces *fos-1a(-)* in the AC (normal AC invasion versus block 7/30) (B) Representative images of FLP-ON^lin-29^::TIR1 system, AC (*cdh-3p*::mCh::moeABD), BM and mNG::mIAA7::FOS-1A. 5-Ph-IAA induces *fos-1a(-)* in the AC (normal AC invasion versus block 24/38). Open arrowheads indicate ACs. Yellow arrowheads indicate boundary of the breach on the BM. White brackets indicates 1° VPC. (C) Schematic overview of IAA proteins and the different AID degrons that have been derived from them. IAA = Indole-3-acetic acid; AID = auxin inducible degron; I = domain I; KR = conserved lysine and arginine residue; II = domain II; PB1 = Phox and Bem1p domain. (D) Quantification of mNG::AID-tagged and mNG::mIAA7-tagged FOS-1A MGV in individual AC nuclei, with FLP-ON^lin-29^::TIR1 mediated FOS-1A depletion (*n*≥30 animals per treatment, *P*<0.0001, statistical comparison was made by Student’s *t*-test between 5-Ph-IAA treated animals and control). (E) Percentage of AC invasion defects when loss of FOS-1A in the AC using different degrons (AID and mIAA7). *n*≥30 animals per treatment, statistical comparison was made by Fisher’s exact probability test between 5-Ph-IAA treated animals and control.

We hypothesized there could be several potential reasons for this insufficient AC-specific FOS-1A degradation. The *lin-29* promoter initiates expression in the AC shortly after the AC/VU decision, around the time of the L2/L3 transition. Almost immediately after AC cell fate specification, the AC upregulates pro-invasive TFs such as *fos-1a* and begins polarization of the F-actin cytoskeleton (Lohmer et al. 2014). Thus, there is a short time window (~4-6 hours) for FLP to excise the >STOP> cassette, transcribe _At_TIR1(F79G)::T2A::DHB::2xmK2, and degrade the mNG::AID::FOS-1A prior to invasion. Second, although FLP-mediated recombination will bring _At_TIR1(F79G)::T2A::DHB::2xmK2 under control of the strong *rpl-28* promoter, the pool of _At_TIR1(F79G) may not be sufficient to achieve complete degradation in the short window of time between genomic excision and AC invasion. In support of this, we quantified the mean intensity of DHB::2xmK2 as proxy for _At_TIR1(F79G) levels (Ahier and Jarriault 2014) between ubiquitous, somatic gonad, and AC-specific FLP-excised lines. At the mid-L3 stage, we found no significant difference in mean DHB::2xmK2 levels between *ckb-3p*::FLP excised _At_TIR1(F79G)::T2A::DHB::2xmK2 and ubiquitous expression of the *rpl-28p* driven transgene. However, *lin-29p*::FLP was only able to generate 12.5% of _At_TIR1(F79G)::T2A::DHB::2xmK2 in the AC at the normal time of invasion (Fig. S5 A-B). These results suggest that levels of _At_TIR1 may still be limiting following AC-specific recombination. Given the narrow window of time needed for rapid degradation of a target protein in the AC to perturb invasion and the dependence on transcription in our FLP-ON^lin-29^::TIR1 system following excision, we measured levels of FOS-1A in early L4 stage animals at the P6.p 8-cell stage, ~4-6 hours later, with animals showing near complete AC-specific depletion of mNG::AID::FOS-1A (Fig. S6). At this time, even with undetectable FOS-1A, we observed no AC invasion defects (30/30 animals). This is not unexpected, however, as invasion is initiated at the P6.p 2-cell stage, when live imaging has detected that a single F-actin-rich protrusion breaches the BM (Hagedorn et al., 2013). Thus, to block AC invasion, it is necessary to accelerate the degradation kinetics of the system to degrade our target protein more rapidly.

A recent report in *C. elegans* demonstrated that an alternative degron sequence, the mIAA7 degron, displayed faster degradation kinetics when paired with _At_TIR1 expression and auxin treatment (Sepers et al. 2022). Compared to the minimal AID epitope, the mIAA7 sequence contains the 44aa domain required for _At_TIR1 recognition and additional N-terminal flanking sequences from the IAA protein that improve the interactions with TIR1 (Gray et al. 2001; Ramos et al. 2001) (Fig. 4C). To test this new degron, we used CRISPR/Cas9 genome engineering to generate an mNG::mIAA7::FOS-1A allele to pair with our FLP-ON^lin-29^::TIR1 system. Additionally, in this iteration, we also used an AC-specific reporter of the F-actin cytoskeleton (*cdh*-*3p*::mCh::moeABD), to facilitate scoring of AC invasion (Matus et al. 2015). Strikingly, compared to AID-tagged FOS-1A, mIAA7-mediated degradation of FOS-1A achieved a more penetrant AC invasion defect (24/38) with more FOS-1A protein eliminated in the AC (70%, Fig. 4B, D, E). Additionally, we observed that FOS-1A degradation was explicitly limited to the AC (Fig. 4B, merged channel), leaving FOS-1A expression in surrounding VU cells (Fig. S7). Importantly, this system achieved AC-specific BM invasion phenotypes without eliciting FOS-1A-dependent cell-cycle arrest phenotypes in adjacent uterine cells as described above (Fig. S7).

Through examination of the AC-specific F-actin reporter, we also observed that degradation of FOS-1A resulted in non-invasive ACs generating mislocalized F-actin rich protrusions apicolaterally instead of along the basal invasive membrane compared to control animals (Fig. S8). While *fos-1a(-)* ACs have been shown to still generate invadopodia (Lohmer et al. 2014), this mispolarized protrusive phenotype suggests an underlying connection between the gene regulatory network controlled by FOS-1A, the F-actin cytoskeleton and polarity regulation mechanisms in the invasive AC.

### AC specific degradation of abundant structural proteins is more efficient by an alternative degron sequence, mIAA7

Although mIAA7-dependent depletion of FOS-1A did not completely recapitulate the null allele phenotype, it significantly improved both the degradation kinetics and associated invasion defects following loss of a transcription factor with single cell specificity. This is in support of the first report using the mIAA7 degron in *C. elegans*, which emphasized that nuclear localized targets are more resistant to auxin-induced degradation (Sepers et al. 2022). To further explore how protein abundance and subcellular localization of a target protein may affect degradation kinetics at single-cell resolution in a functional assay, we generated two new strains to compare the mIAA7 and AID degron kinetics following depletion of *arx-2*, the sole Arp2 subunit of the actin-related protein-2/3 (Arp2/3) complex in *C. elegans* (Sawa et al. 2003). The Arp2/3 complex is an abundant (Wang et al. 2015) branched F-actin regulator and has been a central player in models of protrusive force production via the dynamic actin network (Goley and Welch 2006; Swaney and Li 2016). Using RNAi and an orthogonal dominant negative approach, ARX-2 was previously shown to regulate the polarization of the F-actin cytoskeleton during AC invasion (Caceres et al. 2018). The stability and abundance of the Arp2/3 complex, however, is known to reduce the effectiveness of RNAi in targeting the complex for degradation (Wu et al. 2012 and Zhu et al. 2016), making Arp2/3 an ideal target to test our AC-specific degradation strategy. Following 5-Ph-IAA treatment, both AID-and mIAA7-tagged ARX-2 are eliminated in the AC (Fig. 5A-B), with similar penetrance in AC invasion defects (AID, 52% (19/38); mIAA7, 53% (20/37)) (Fig. 5C). Concomitant with a defect in invasion, we observed that the F-actin cytoskeleton, which is normally polarized at the basal invasive membrane was ectopically enriched apically or laterally in ARX-2-depleted ACs (Figure 5A-B, S9). Notably, while the F-actin enrichment was significantly reduced in both tagged *arx-2* alleles, we observed a greater polarity defect in the mIAA7-tagged allele, with a ~2-fold reduction in mean polarity value with the mIAA7 degron as compared to a ~1.3 fold change with an AID degron (Fig. 5D). Together, our results suggest that the mIAA7 degron performs better than AID for nuclear localized targets or even other subcellular compartments within a lineage. This is particularly important when speed and degree of degradation is a critical component of experimental workflow.

**Figure 5.**
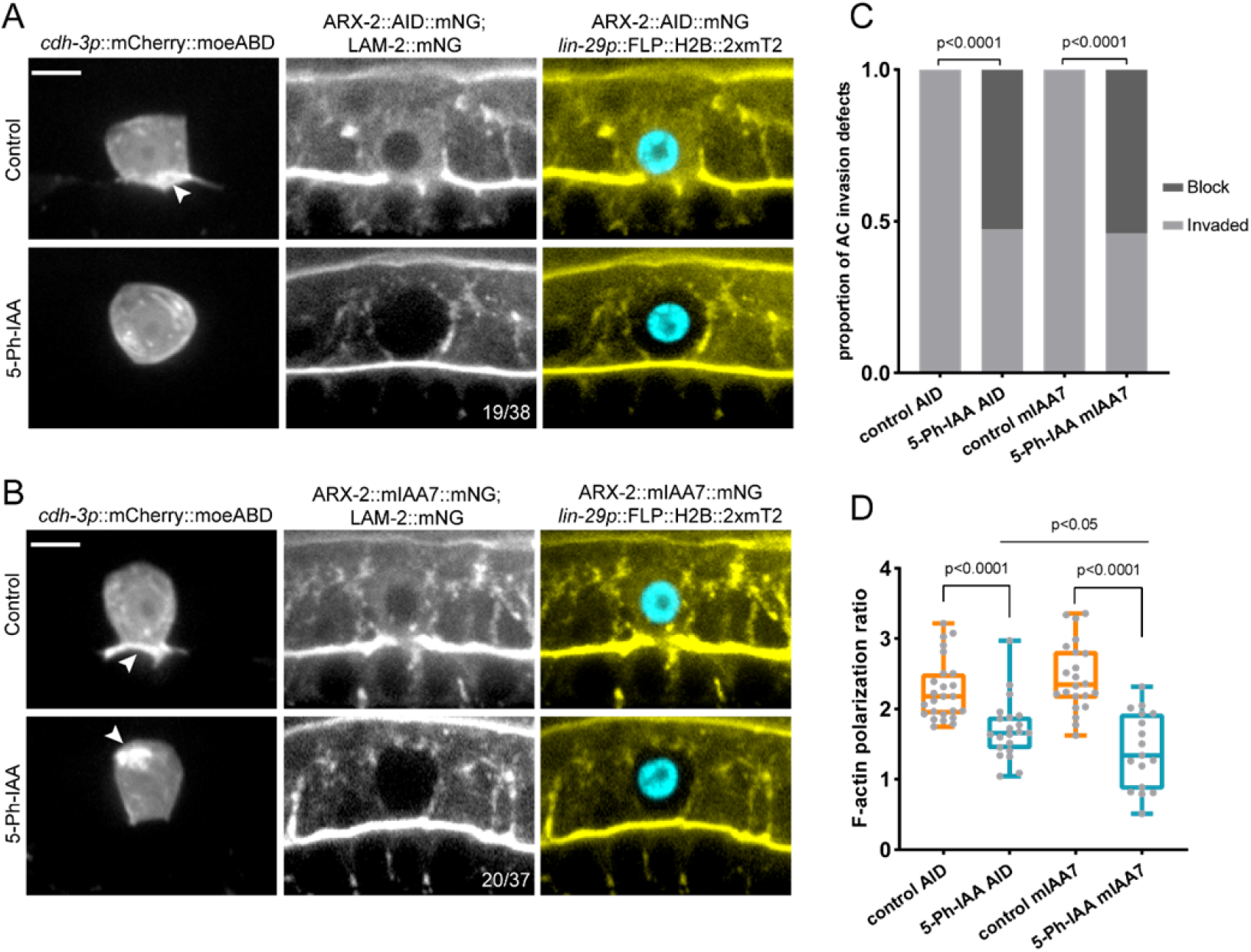
AC specific degradation of ARX-2 using AID and mIAA7 degrons. (A-B) Micrographs depicting AC specific degradation of ARX-2::mIAA7::mNG (A) and ARX-2::AID::mNG using FLP-ON^lin-29^::TIR1. Left channel shows AC F-actin (*cdh-3p*::mCh::moeABD) polarization, middle channel shows BM and endogenously mIAA7/AID tagged ARX-2, right channel overlayed with *lin-29p*::FLP::H2B::2xmT2. (C) Percentage of AC invasion defects when loss of ARX-2 in the AC using mIAA7 and AID. *n*≥30 animals per treatment, statistical comparison was made by Fisher’s exact probability test between 5-Ph-IAA treated animals and control. (D) F-actin polarization ratio is calculated via mean intensity of the F-actin at invasive membrane divided by apicolateral of ACs. *n*≥20 animals per treatment, statistical comparison was made by Wilcoxon test between 5-Ph-IAA treated animals and control. Mann-Whitney test was used to compare normalized ratio of 5-Ph-IAA AID and 5-Ph-IAA mIAA7, p=0.0167, significant at p<0.05. Scale bars: 5 μm.

### An expandable FLP-ON::TIR1 system for multifunctional applications

In the previous experiments we demonstrate how the FLP-ON::TIR1 system can be combined with a fluorescent biosensor for simultaneous readout of protein degradation and cell-cycle state. To extend the utility of the system to the broader *C. elegans* community, we have generated a series of adaptations. First, we generated two additional tissue/cell specific FLP drivers. The first uses the *rgef-1* promoter (Zeynep et al. 2001) to generate _AT_TIR1(F79G) in neurons and the second utilizes the *wrt-2* promoter (Gudrun et al. 1999) to drive _AT_TIR1(F79G) in hypodermal cells (Fig. 6B). Second, we introduced membrane localization sequences in the FLP-ON::TIR1 construct. Specifically, we added a 10 amino-acid N-terminal dual acylation motif of LCK (Zlatkine et al. 1997) fused to mNG internal to the FRT_3_ sites of the STOP cassette (>LCK::mNG::STOP*>)* generating a strain with ubiquitous bright green membrane localization prior to recombination. We also swapped the CDK sensor for a second membrane localization sequence, using the PLC-δ-PH domain (Hurley 2006) which can also target fusion proteins to the plasma membrane (_At_TIR1(F79G)::T2A::PH::2xmK2). We generated this new construct with both the *rpl-28* promoter as well as a ccdB cassette (Philippe et al. 1994) to enable flexible promoter swapping. With these adaptations, depleting new degron-tagged knock-in alleles in a tissue or cell type of interest through the FLP-ON::TIR1 system can be done using standard genetic crosses or microinjections (Dokshin et al. 2018)) (Fig. 6A). To demonstrate the efficacy of these adaptations, we crossed the pan-neuronal (*rgef-1*) and seam cell (*wrt-2*) FLP strains into the dual FLP-ON::TIR1 reporter. As expected, we detected bright green membrane localization in all cells and strong red membrane localization in cells co-expressing FLP (Fig 6C). Thus, the FLP-ON::TIR1 system, combined with the set of FLP drivers described here and a growing list of tissue-specific FLP drivers generated by the community, can be used to explore novel gene functions in a cell type or lineage-restricted fashion.

**Figure 6.**
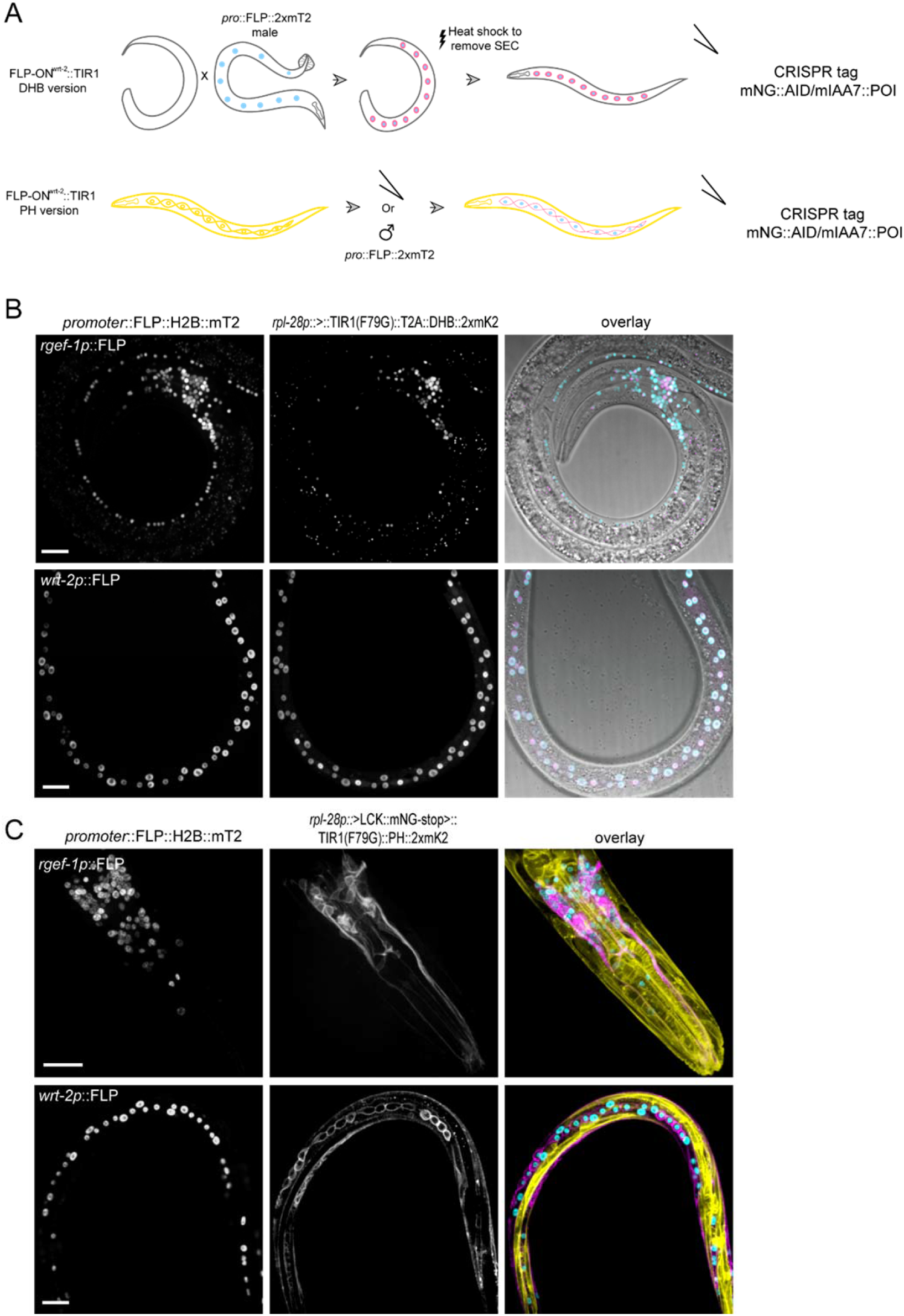
An expandable FLP-ON::TIR1 system. (A) Workflow and examples using a hypodermal-driven FLP (FLP-ON^wrt-2^::TIR1) for strain construction using standard genetic crosses or microinjection to pair the FLP-ON::TIR1 system with a degron tagged protein of interest (POI). (B) *rgef-1p*::FLP and *wrt-2p::FLP* with *rpl-28p*::>STOP>::_At_TIR1(F79G)::T2A::DHB::2xmK2, demonstrating FLP::H2B::2xmT2 expression (left panel), _At_TIR1(F79G)::T2A::DHB::2xmK2 expression (middle panel), and overlayed with DIC (right panel). (C) *rgef-1p::FLP* and *wrt-2p::FLP* with *rpl-28p*::>LCK::mNG::STOP>::_At_TIR1(F79G):: T2A::PH::2xmK2, demonstrating FLP::H2B::2xmT2 expression (left panel), _At_TIR1(F79G):: T2A::DHB::2xmK2 expression (middle panel), and overlayed with rpl-28p::>LCK::mNG, yellow (right panel). Scale bars, 5 μm.

## Discussion

In specific contexts, tissue-or cell type-specific expression of heterologous _At_TIR1 using spatially restricted promoters is insufficient to degrade AID-tagged target proteins. This is likely due to either lower levels of transcription from promoters that can drive restricted expression patterns or because many developmentally regulated promoters are dynamically expressed. In either of these cases, the normal degradation kinetics of AID-tagged targets are slower than those observed when _At_TIR1 is expressed from ubiquitously expressed promoters (e.g., *eft-3* or *rpl-28)*. In this study, we set out to develop a general two-component system that takes advantage of tissue-or cell type-specific promoters to drive restricted FLP activity that can re-animate expression from a dormant, high-powered promoter with cellular resolution. We accomplished this by inserting an FRT_3_::STOP::FRT_3_ (>STOP>) cassette into the well-characterized *rpl-28* promoter that is efficiently expressed in both somatic and germline tissues. We demonstrate that a variety of tissue-specific promoters can be used to reanimate this dormant transgene and that the level of _At_TIR1(F79G) activity derived from the FLP-ON::TIR1 dramatically improves AID-target degradation. Given the recombination event is permanent and the induced _At_TIR1(F79G) is not active until the addition of 5-Ph-IAA (Hills-Muckey et al. 2022; Negishi et al. 2022), the system combines a standardized approach to the spatial and temporal depletion of any degron-tagged target protein.

We use the FLP-ON::TIR1 system to reveal genetically separable, cell type-specific phenotypes for multiple important regulators of differential cell identity. First, we use this system to demonstrate cell specific functions of a classic signaling molecule, LIN-12/Notch, during cell fate specification in the somatic gonad and developing vulva. We separately deplete LIN-12 in the developing VPCs and then in the somatic gonad to demonstrate that some VPC fate transformations that likely occur in genetic *lin-12(0)* animals are potentially derived from both the misspecification of VU cells and the ectopic secretion of the inducing LIN-3 ligand from supernumerary ACs. We also use the FLP-ON::TIR1 system to identify novel phenotypes associated with depleting the conserved FOS-1A transcription factor implicated in differential transcriptional programs that control the cellular behaviors of the invasive AC and proliferative uterine cells. Specifically, we identify a new role for FOS-1A in maintaining the proliferative state of VU cells during the L3 stage. Finally, we demonstrate that the FLP-ON::TIR1 system is compatible with the mIAA7 AID epitope and that this leads to an even more efficient tissue-specific degradation of nuclear factors as well as a cytoskeleton protein, ARX-2, a core component of the Arp2/3 complex.

While we used this system to characterize phenotypes associates with depleting expression in the somatic gonad or in developing VPCs, we envision that this system will be useful to dissect the roles of other developmental genes as well as many genes that are essential for development but exhibit lethal phenotypes at the organismal level. For example, many of factors involved in chromatin remodeling (van der et al. 2020), mRNA processing/splicing (Arribere et al. 2020), protein post-translational regulation (Wenzel et al. 2011) and miRNA processing/function work differentially in diverse cell types to control separate gene regulatory networks (Kato and Slack 2008). The FLP-ON::TIR1 system can be employed to identify these differentially regulated targets and determine the phenotypes associated by specifically inhibiting these processes in individual cells. This combined with the temporal control of the _At_TIR1 system (through timed addition of 5-Ph-IAA) makes the tractability of this system immense.

Finally, we generated an expandable toolkit using the basic FLP-ON::TIR1 system that can be further modified by other *C. elegans* researchers. The modular nature of the system allows for two features to be independently modified. First, any promoter can be used to drive specific expression of a FLP construct. With a growing list of promoters that have been characterized by many researchers worldwide, paired with previous genome-wide promoter activity analyses in transgenic *C. elegans* strains (Dupuy et al. 2007), a wide range of AID-tagged proteins could be rapidly depleted in defined cell types. In cases where an characterized individual promoters are not specific enough to drive expression in a defined cell type, precise spatial degradation could be achieved by overlapping the expression region of FLP and _At_TIR1, as has been done to achieve neuronal intersectional labeling through a split Q system (Wei et al. 2012). Second, we demonstrate the utility of co-expressing a variety of other fluorescently tagged reporters that can be efficiently co-expressed with TIR1 in targeted tissues. These reporters can be inserted immediately after the 2A peptide sequence in the _At_TIR1(F79G)::T2A construct and allow any type of reporter transgene to be co-expressed from the reactivated promoter. We demonstrate the utility of this feature using our cell-cycle sensor based on CDK activity, as well as a dual membrane (LCK/PH) reporter. We can predict several interesting applications to expand the employment of this 2A-based co-expression system. For instance, for cell biological assays, diverse morphologies of the actin cytoskeleton could be monitored by LifeAct across tissues (Garcia et al. 2022). For examining metabolic phenotypes, a genetically encoded Förster resonance energy transfer (FRET)-based ATP biosensor (ATeam) (Tsuyama et al. 2013) or mitochondrial Ca^2+^ sensor (Alvarez-Illera et al. 2017) could be co-transcribed when metabolic signaling molecules are perturbed. For probing questions in developmental biology or aging, gene regulatory networks could be explored through the degradation of a targeted transcription factor with simultaneous misexpression of downstream effectors. Finally, to gain insight into protein complexes containing multiple subunits, one could co-express dominant negative orthologs while targeting the degradation of other components to further eliminate complex function within a lineage or cell type. Together, this system should be feasible in any context amenable to recombination-based methods and applicable to diverse biological areas including cell and developmental biology, neuroscience, immunology, metabolism and aging research.

## Materials and methods

### Plasmid construction

Throughout this study, we used conventional restriction enzyme-mediated cloning and Gibson assembly technology. The primers used in this study as well as promoter lengths are described in Table S1 in the Supplemental Material.

To generate plasmid pYX026_ccdb::FLP, we digested pWZ159 with ClaI/AflII as the backbone, and PCR amplified FLP from pYX014 with YX42/YX43 for Gibson assembly. Plasmids pYX027_*ckb-3p*::FLP, pYX029_*egl-43p*::FLP, *pYX033_unc-62p*::FLP, were made through Gibson assembly by replacing the ccdb site with 2 kb of the *ckb-3* promoter, 1.7 kb *egl-43* promoter, 3.4kb *unc-62* promoter. Plasmid containing histone H2B tagged with 2xmT2 (pYX028_*ckb-3p*::FLP::P2A::H2B::2xmT2) was designed to provide visualization of FLP expression. Fragments P2A::H2B, and H2B::2xmT2 were PCR amplified from pTNM114 with YX46/YX47, pYX014 with YX48/YX49, respectively. Plasmids pYX030_*lin-29p*::FLP::P2A::H2B::2xmT2, pYX038_*rgef-1p*::FLP::P2A::H2B::2xmT2, and pYX039_*wrt-2p*::FLP::P2A::H2B:: 2xmT2 were constructed using pYX028 as backbone and the *ckb-3p* fragment was replaced with *lin-29p*, *rgef-1p*, *wrt-2p*, respectively.

Plasmid pYX023_*rpl-28p*::>STOP>::_At_TIR1(F79G)::T2A::DHB::2xmK2 contains the _At_TIR1(F79G) coding sequencing and a fragment of human DNA helicase B (DHB) fused with 2xmKate2 (Hills-Muckey et al. 2022; Martinez et al. 2022) (STOP cassette flanked by FRT3 sites includes two stop codons, TAATAG, followed by the *let-858*-3’UTR).

### C. elegans strains and culture conditions

Animals were maintained under standard conditions and cultured at 20-25°*C* on NGM plates. Animals were synchronized for experiments through alkaline hypochlorite treatment of gravid adults to isolate eggs (Porta-de-la-Riva et al. 2012). In the text and figures, we designate linkage to a promoter through the use of a (*p*) and fusion of a proteins via a (::) annotation. A complete list of strains can be found in Table S2.

### Molecular biology and microinjection

The mNG::AID::FOS-1A allele was generated by the SEC method using CRISPR/Cas9 genome editing via microinjection into the hermaphrodite gonad (Dickinson et al. 2015). Repair templates were generated as synthetic DNAs from either IDT or Twist Biosciences. These synthetic fragments were cloned into ccdB compatible sites in pDD282 by Gibson assembly (New England Biolabs). Homology arms for each targeting construct are about 1kb on 5’ and 3’ to the epitope insertion site (see Tables S3 for additional details). sgRNAs were constructed by EcoRV and NheI digestion of the plasmid pDD122. This was accomplished by amplifying a 230 bp fragments that was used to replace the sgRNA targeting sequence from pDD122 with a new sgRNA using Gibson assembly (New England Biolabs). Hermaphrodite adults were co-injected with guide plasmid (50 ng/μl), repair plasmid (50 ng/μl) and an extrachromosomal array marker (pCFJ90, 2.5 ng/μl). Injected animals and their progeny were incubated at 25°C for three days before carrying out genotyping to identify successfully edited loci. Following identification of CRISPR-mediated insertions, floxing of the SEC was carried out using standardized protocols (Dickinson et al. 2015).

Alleles of *arx-2* and *fos-1a* (ARX-2::mIAA7::mNG, ARX-2::AID::mNG and mNG::mIAA7::FOS-1A) were generated using PCR amplified repair templates, using CRISPR/Cas9 genome editing via microinjection into the hermaphrodite gonad (Dokshin et al. 2018). 1.52mM cas9 protein, 1.52mM tracer RNA, 1.52mM guide RNA, and 0.2-2mM PCR products as repair templates mixed with co-injection marker (*rol-6*) were injected into the worms. PCR products were amplified from plasmid containing mNG::mIAA7::AID::linker (pWZ297), or linker::mIAA7::mNG (pWZ296) from vectors with ultramer oligonucleotides containing 100bp homology sequence.

The guide RNAs and repair templates for each gene are listed in Tables S3.

### Live-cell imaging

All micrographs in this manuscript were collected on a Hamamatsu Orca EM-CCD camera mounted on an upright Zeiss AxioImager A2 with a Borealis-modified CSU10 Yokagawa spinning disk scan head (Nobska Imaging) using 440nm, 505nm, 488 nm and 561 nm Vortran lasers in a VersaLase merge and a Plan-Apochromat 100×/1.4 (NA) or 40x/1.4NA Oil DIC objective. MetaMorph software (Molecular Devices) was used for microscopy automation. Several experiments were scored using epifluorescence visualized on a Zeiss Axiocam MRM camera, also mounted on an upright Zeiss AxioImager A2 and a Plan-Apochromat 100×/1.4 (NA) Oil DIC objective. Animals were mounted into a drop of M9 on a 5% Noble agar pad containing approximately 10 mM sodium azide anesthetic and topped with a coverslip.

### Auxin induced protein degradation

For all auxin experiments, synchronized L1 larval stage animals were first transferred to standard nematode growth media (NGM) agar plates seeded with *E. coli* OP50 and then transferred at the P6.p 2-cell stage (mid-L3 stage) to either OP50-seeded NGM agar plates treated with 0.1mM 5-Ph-IAA, 4mM NAA or 1mM K-NAA. 5-Ph-IAA, NAA and K-NAA were diluted into the NGM agar (cooled to approximately 50°) at the time of pouring plates. Fresh OP50 was used to seed plates. For control experiments, OP50-seeded NGM agar plates containing 0.25% ethanol were used.

### Assessment of AC invasion and specification

Invasion of the AC into the vulval epithelium was scored by the visualization of the CDK activity sensor (Adikes et al., 2020; Martinez et al. 2022) and a gap in the basement membrane underneath the AC. In strains with the endogenous labeled LAM-2::mNG, an intact yellow fluorescent barrier under the AC was used to assess invasion. Wild-type invasion is defined as a breach as wide as the basolateral surface of the AC at the P6.p 4-cell stage (Sherwood and Sternberg 2003). Raw scoring data is available in the supplemental material.

When using FLP-ON^ckb-3^::TIR1 and FLP-ON^lin-29^::TIR1 to deplete mNG::AID::FOS-1A to assay the FLP system efficiency, AC invasion was scored at the P6.p 4-cell stage, when 100% of wild-type animals exhibit a breach in the BM (Sherwood and Sternberg 2003). The “2AC” phenotype was scored at the P6.p 4-cell stage when depleting LIN-12::mNG::AID in the somatic gonad using FLP-ON^ckb-3^::TIR1.

### Image quantification and statistical analysis

Images were processed using Fiji/ImageJ (v.2.1.0/1.53c) (Schindelin et al. 2012). Expression levels of mNG::AID::FOS-1A, DHB::2xmK2 were measured by quantifying the mean gray value of AC nuclei, defined as the somatic gonad cell near the primary vulva arrested in a CDK_low_, G0 state (or with *lin-29p*::FLP::H2B::mT2 expression). Background subtraction was performed by rolling ball background subtraction (size = 50). Quantification of the CDK activity sensor (DHB::2xmKate2) was performed by hand in ImageJ, as previously described (Adikes et al. 2020). AC F-actin polarity was determined using the ratio of the mean fluorescence intensity (Kelley et al. 2019) from a 5-pixel-wide line scan drawn in ImageJ along the invasive and apicolateral membranes of ACs. Polarity was calculated as the following ratio: [invasive membrane mean intensity- background] / [apicolateral membrane mean intensity-background].

Images were overlaid and figures were assembled using Adobe Photoshop 2020 and Adobe Illustrator 2020, respectively. Statistical analyses and plotting of data were conducted using GraphPad Prism (v. 8.0.2). Statistical significance was determined using either an unpaired two-tailed Student’s *t*-test or Fisher’s exact probability test. Figure legends specify when each test was used and the p-value cut-off.

## Data availability

Strains and plasmids are available upon request. The authors affirm that all data necessary for confirming the conclusions of the article are present within the article, figures, and tables.

Supplementary material is available at GENETICS online.

## Acknowledgments

We would like to thank members of the Hammell, Shen and Matus laboratories for the critical review of this manuscript. We received strains from Peter Askjaer and the Caenorhabditis Genetics Center (CGC), which is funded by the NIH Office of Research Infrastructure Programs (P40 OD010440). We would like to thank Peter Askjaer for the initial suggestions for building FLP constructs. We thank T.N.M. Kinney and J. Smith for providing additional feedback.

---

**Figure S1.**
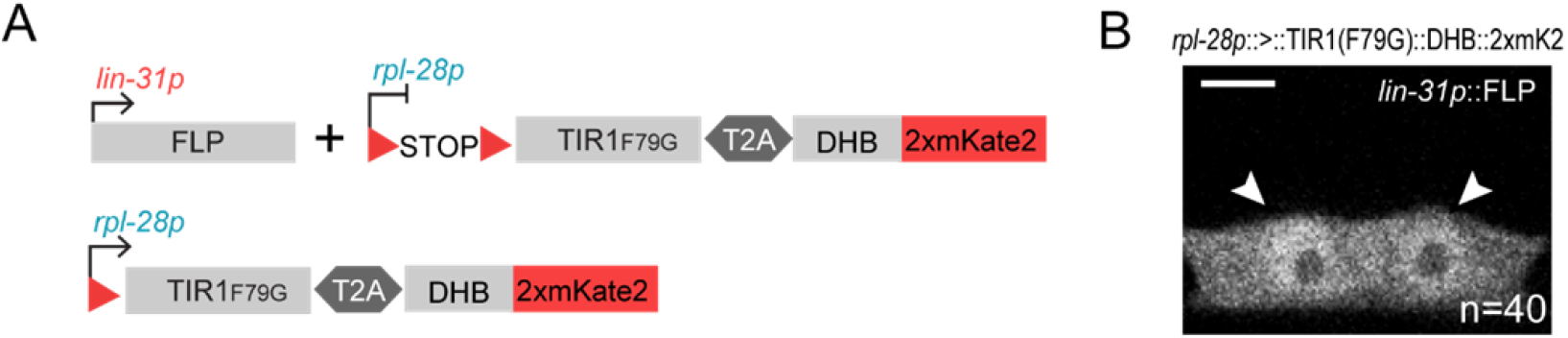
*lin-31p*::FLP efficiently restricts *rpl-28p*::>::TIR1(F79G)::T2A::DHB::2xmK2 to the VPCs. (A) Schematic figure depicting *lin-31p::FLP* and *rpl-28p*::>STOP>::_At_TIR1(F79G)::T2A::DHB::2xmK2 constructs. (B) *rpl-28p*::>::_At_TIR1(F79G)::DHB::2xmK2 is specifically expressed in the VPCs, mediated by *lin-31p::FLP*. Scale bar: 5 μm

**Figure S2.**
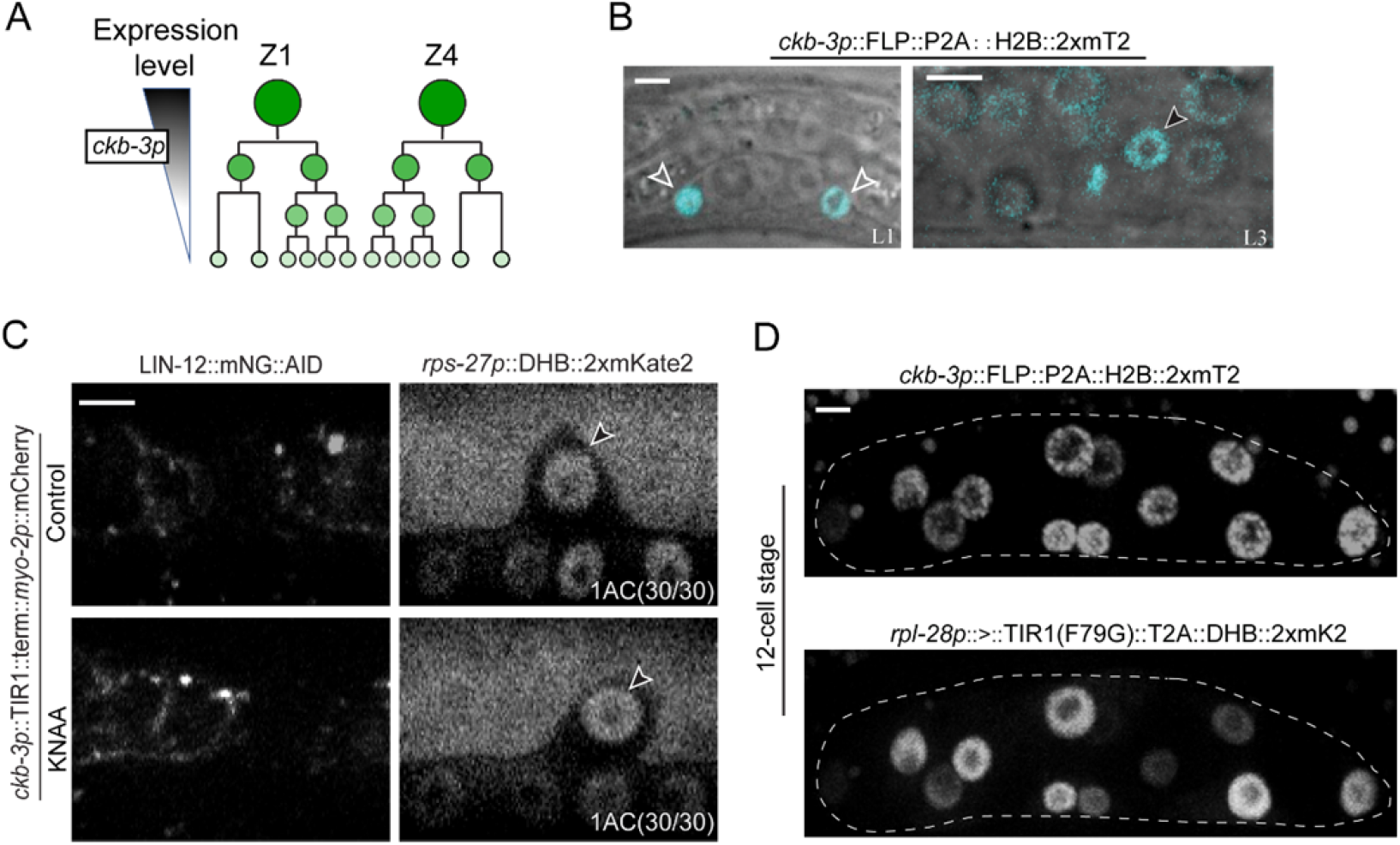
*ckb-3p*::FLP expression pattern and uterine LIN-12/Notch depletion using a *ckb-3p*::TIR1 construct. (A) Schematic of expression pattern and levels of *ckb-3* promoter over developmental time. (B) *ckb-3p*::FLP::H2B::2xmT2 expresses in the Z1/Z4 and their descendants (C) Uterine specific degradation of LIN-12::mNG::AID using *ckb-3p::TIR1::term::myo-2p::mCh* with 1 mM K-NAA treatment. Arrowheads indicate ACs. (D) *ckb-3p*::FLP::H2B::2xmT2 and *rpl-28p*::>::_At_TIR1(F79G)::T2A::DHB::2xmK2 are expressed in the 12 somatic cells derived from Z1/Z4 expression. Scale bars: 5 μm.

**Figure S3.**
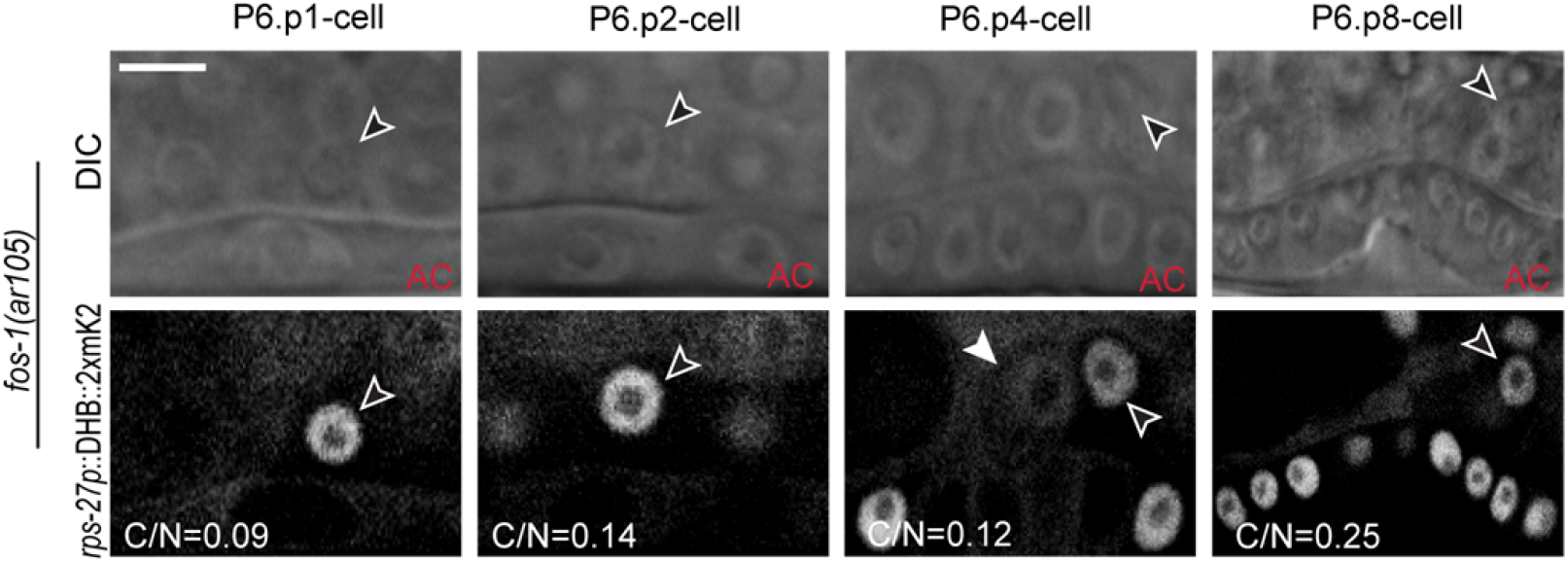
AC invasion defects in *fos-1(ar105)* animals. Representative images (DIC, top and fluorescence, bottom) of the focal plane containing the AC (open arrowheads) in *fos-1(ar105)* animals expressing *rps-27p:*:DHB::2xmK2, during the L3 stage. Cytoplasmic:Nuclear (C/N) ratio-metric quantification of ACs indicates that they are in a CDK_low_, G0-arrested state. VU cells indicated by an solid white arrowhead. Scale bar: 5 μm

**Figure S4.**
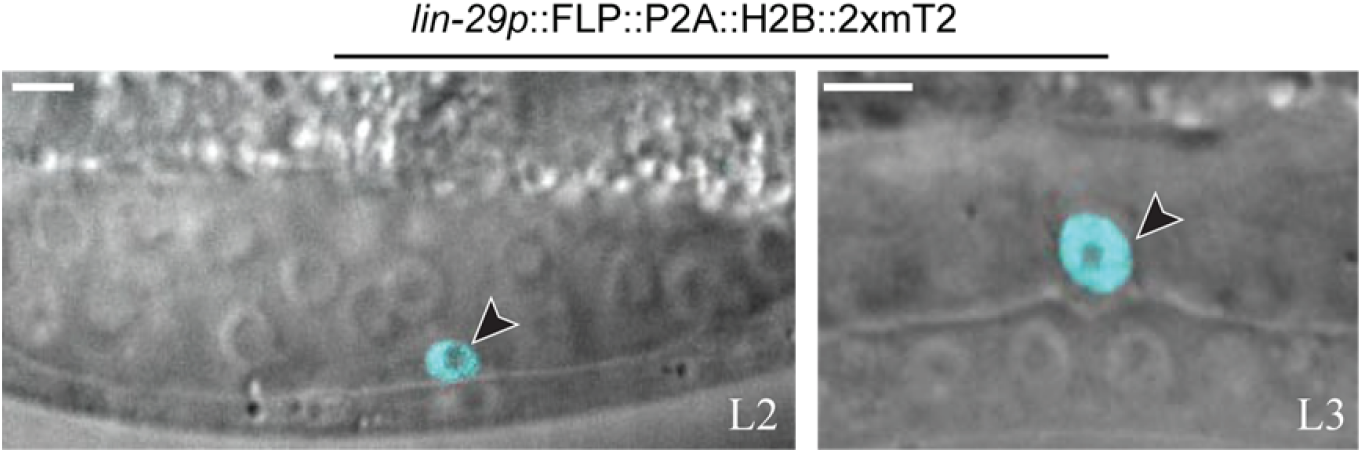
*lin-29p*::FLP::P2A::H2B::2xmT2 expression pattern. Fluorescence overlay of DIC images of representative micrographs depicting *lin-29p*::FLP::P2A::H2B::2xmT2 expression in the specified AC (black arrowheads) during late L2 (left) and middle L3 at normal time of AC invasion (right). scale bar: 5 μm.

**Figure S5.**
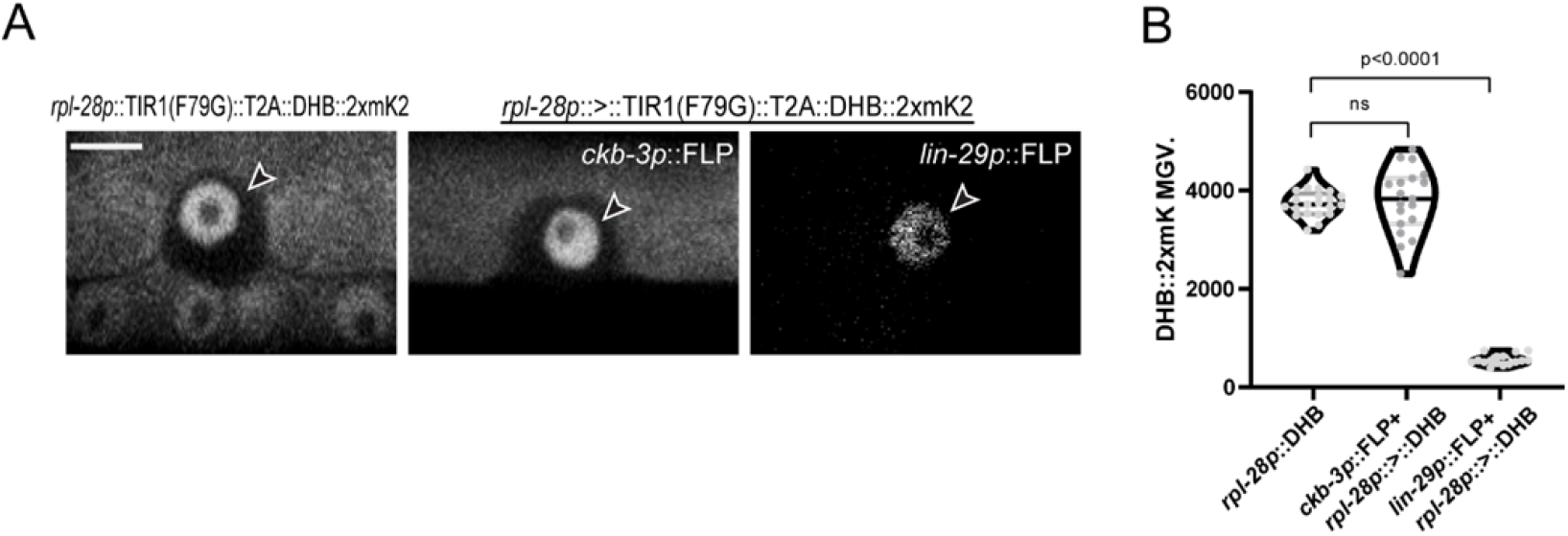
Comparison of DHB intensity. (A) Representative micrographs showing nuclear DHB signal in the AC (top: *rpl-28p*::_At_TIR1(F79G)::DHB::2xmK2; middle: FLP-ON^ckb-3^::TIR1; bottom: FLP-ON^lin-29^::TIR1). Arrowheads indicate ACs, Scale bar: 5 μm. (B) Quantification of DHB::2xmK2 MGV in the AC.

**Figure S6.**
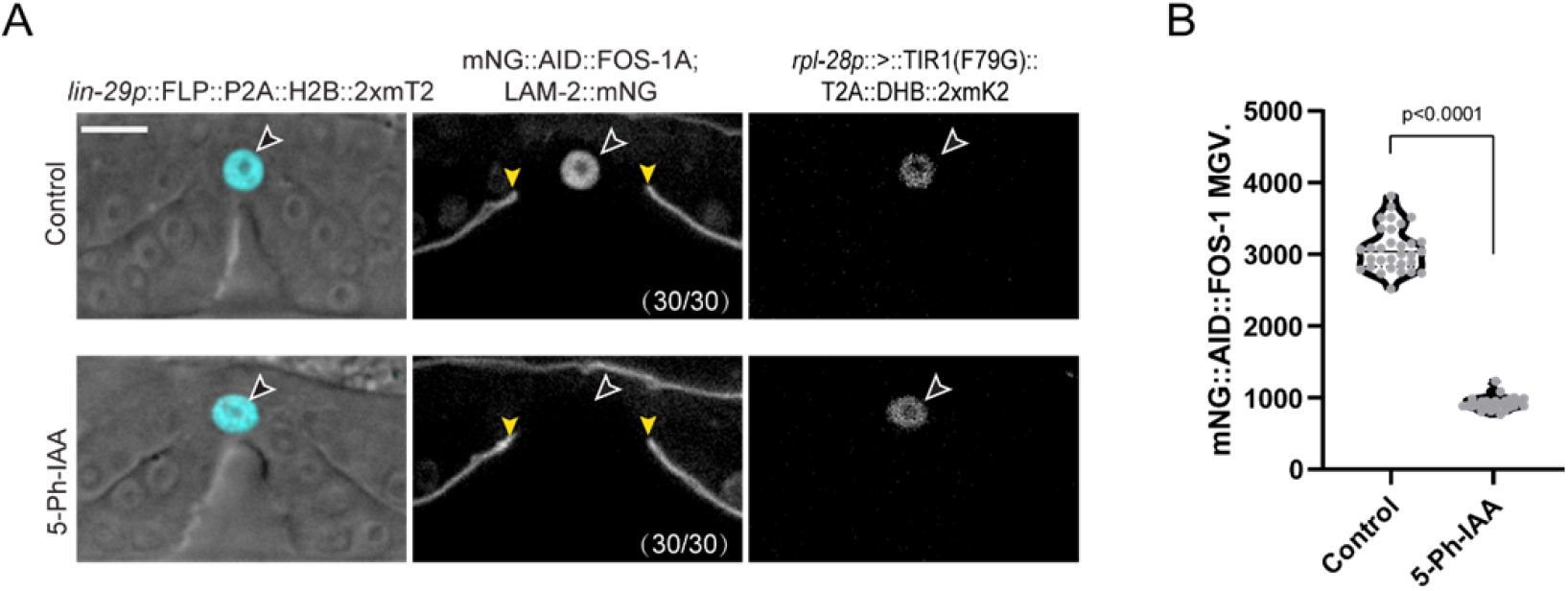
AC specific FOS-1A depletion at P6.p 8-cell stage. (A) Representative micrograph showing FOS-1A degradation in the AC (black arrowheads) by FLP-ON^lin-29^::TIR1, n=30. Yellow arrowheads indicate the boundaries of the BM gap (LAM-2::mNG). Scale bar: 5 μm. (B) Quantification of mNG::AID::FOS-1A MGV of individual AC nuclei, with FLP-ON^lin-29^::TIR1 mediated depletion at the P6.p 8-cell stage. (*n*≥30 animals per treatment, *P*<0.0001, statistical comparison was made by Student’s *t*-test between 5-Ph-IAA treated animals and control).

**Figure S7.**
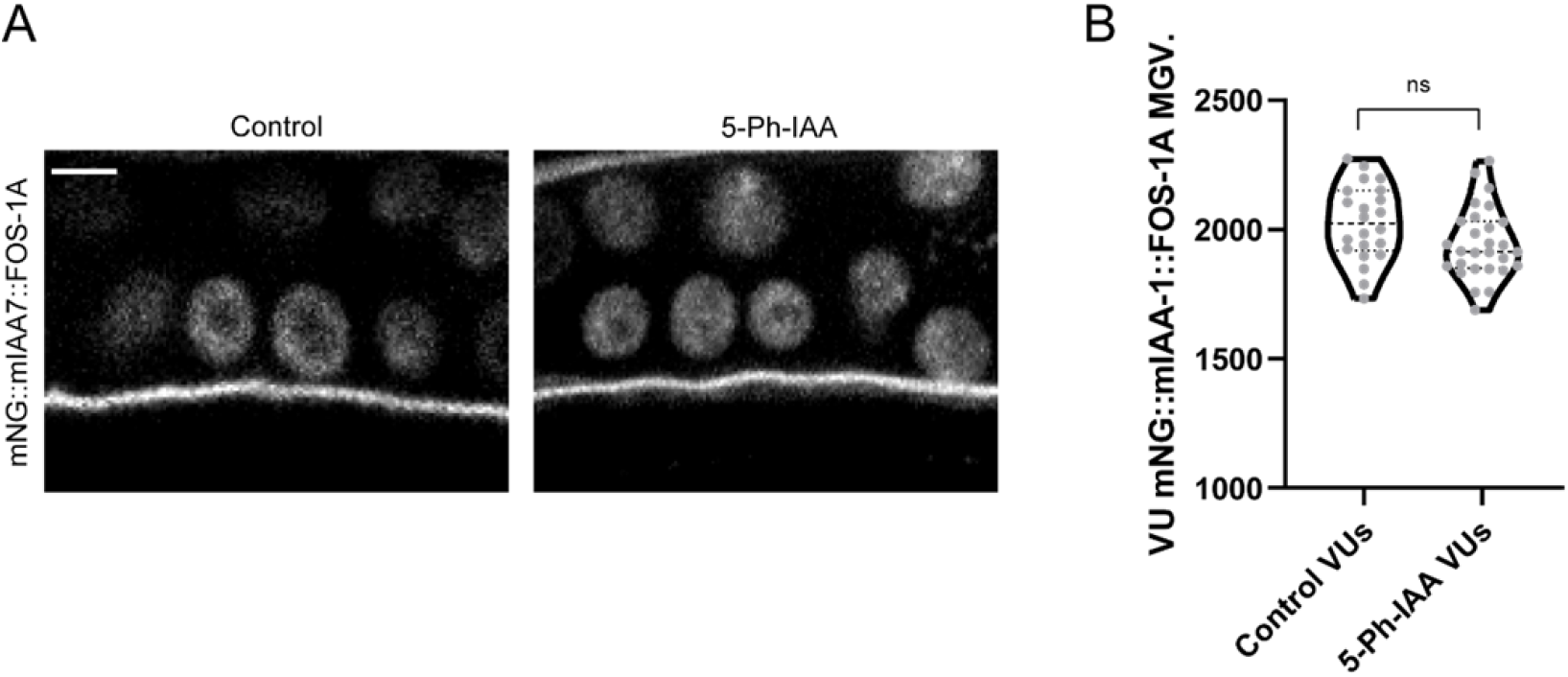
mNG::mIAA7::FOS-1A expression level is not affected in the VU lineage with FLP-ON^lin-29^::TIR1 after auxin exposure. (A) Micrographs depicting mNG::mIAA7::FOS-1A expression in VU cells with and without auxin treatment. (B) Quantification of mNG::mIAA7::FOS-1A MGV in the VU lineage. statistical comparison was made by Student’s *t*-test between control and 5-Ph-IAA treated animals. *n*≥30 animals. ns. Not significant. Scale bar: 5 μm.

**Figure S8.**
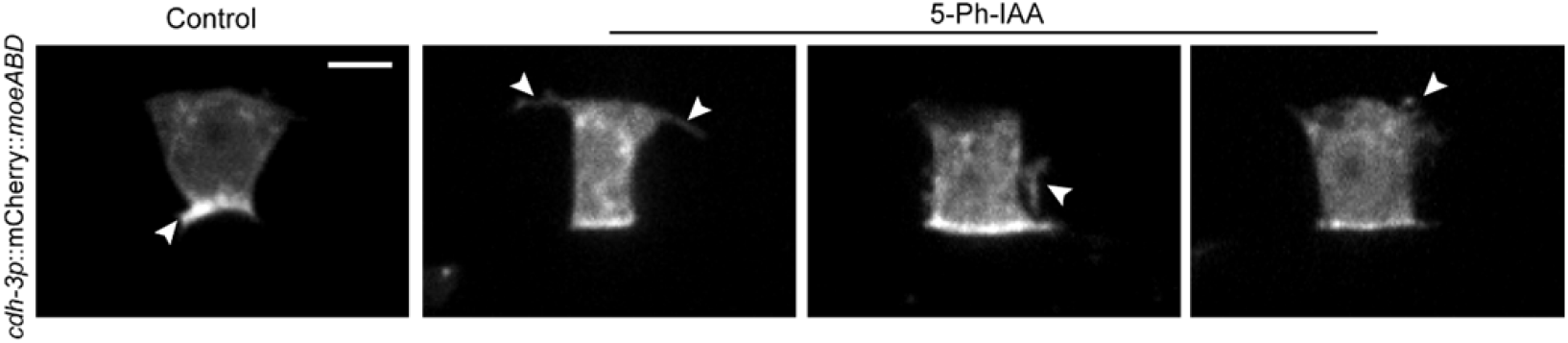
Micrographs depicting representative F-actin (*cdh-3p*::mCh::moeABD) polarization in control animals and FLP-ON^lin-29^::TIR1 mediated mNG::mIAA7::FOS-1A depleted animals. White arrow heads indicate F-actin polarization at invasive membrane (control) and apical-lateral membrane (5-Ph-IAA). Scale bar: 5 μm.

**Figure S9.**
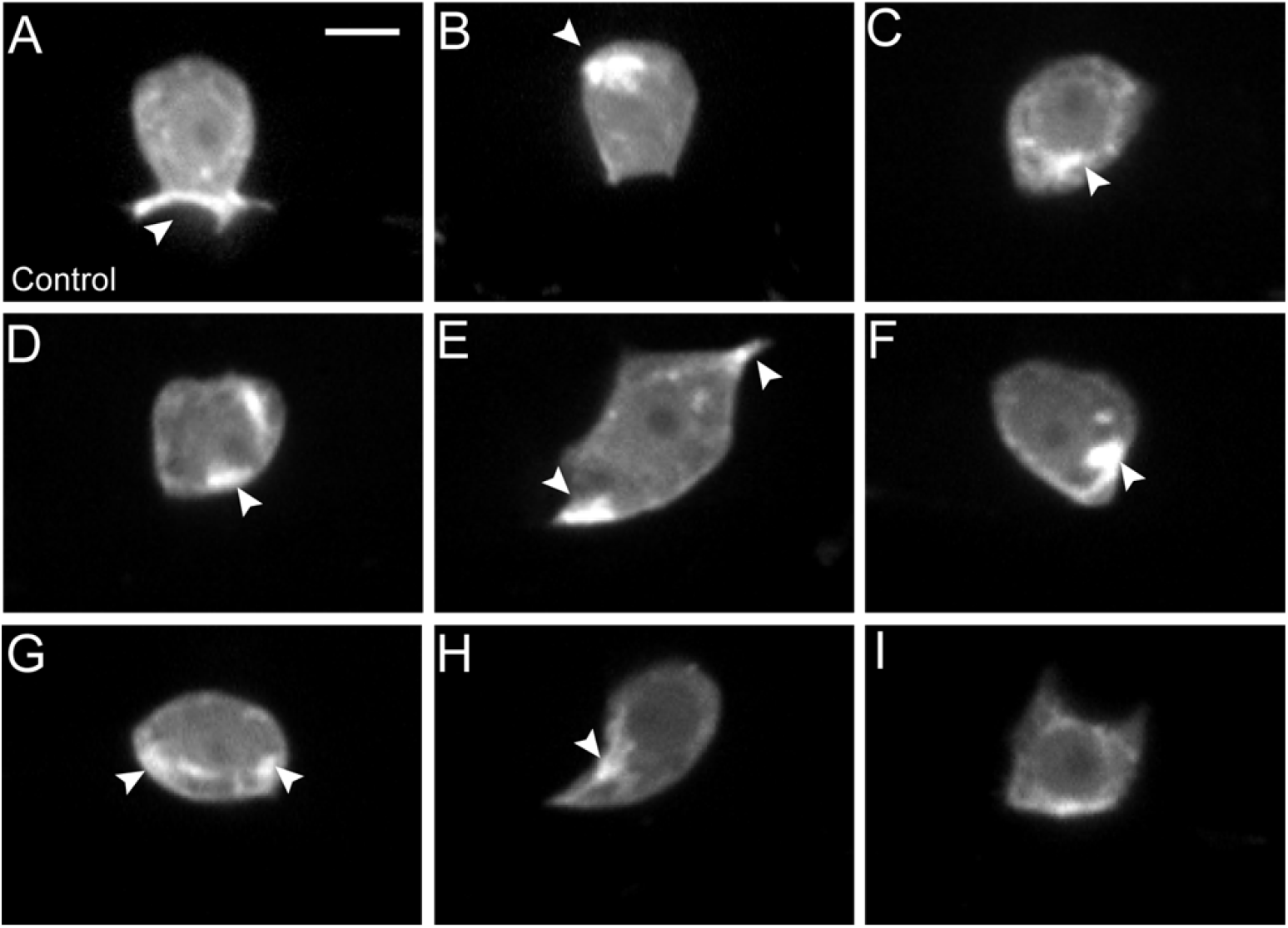
Micrographs depicting representative F-actin (*cdh-3p*::mCh:moeABD) polarization in control animals (A) and FLP-ON^lin-29^::TIR1 mediated ARX-2::mIAA7::mNG degraded animals (B-I). White arrow heads indicate normal F-actin polarization at invasive membrane (control) and mis-localized at apical and lateral membrane (5-Ph-IAA). Scale bar: 5μm.

